# Joint Modeling of Longitudinal Biomarker and Survival Outcomes with the Presence of Competing Risk in Nested Case-Control Studies with Application to the TEDDY Microbiome Dataset

**DOI:** 10.1101/2025.05.23.655653

**Authors:** Yanan Zhao, Ting-Fang Lee, Boyan Zhou, Chan Wang, Ann Marie Schmidt, Mengling Liu, Huilin Li, Jiyuan Hu

## Abstract

**Motivation:** Large-scale prospective cohort studies collect longitudinal biospecimens alongside time-to-event outcomes to investigate biomarker dynamics in relation to disease risk. The nested case-control (NCC) design provides a cost-effective alternative to full cohort biomarker studies while preserving statistical efficiency. Despite advances in joint modeling for longitudinal and time-to-event outcomes, few approaches address the unique challenges posed by NCC sampling, non-normally distributed biomarkers, and competing survival outcomes.

**Results:** Motivated by the TEDDY study, we propose “JM-NCC”, a joint modeling framework designed for NCC studies with competing events. It integrates a generalized linear mixed-effects model for potentially non-normally distributed biomarkers with a cause-specific hazard model for competing risks. Two estimation methods are developed. fJM-NCC leverages NCC sub-cohort longitudinal biomarker data and full cohort survival and clinical metadata, while wJM-NCC uses only NCC sub-cohort data. Both simulation studies and an application to TEDDY microbiome dataset demonstrate the robustness and efficiency of the proposed methods.

## 1 Introduction

The human microbiota is a dynamic community of microorganisms that inhabit various body niches, including the oral cavity, colon, and skin (Ursell 2012). Alterations in microbiota composition can significantly impact health, potentially predisposing individuals to immunological and pathological conditions (Eckburg 2005, Peterson 2009, Blaser 2010). Advances in next-generation sequencing technologies, such as 16S rRNA and shotgun metagenomic sequencing, have facilitated in-depth exploration of the human microbiome’s role in diseases, including Type 1 diabetes (T1D) (Vatanen 2018, Paun 2017, Needell 2016, Dedrick 2020), inflammatory bowel disease (Zhang 2022, Lloyd-Price 2019, Fornelos 2020), and cancer (Rajagopala 2017, Jiang 2021, Jeong 2024).

Large-scale cohort studies, such as the Integrative Human Microbiome Project (iHMP) (Integrative, H.M.P 2019) and the Environmental Determinants of Diabetes in the Young (TEDDY) biomarker study, where gut microbiome is one of the key biomarker profiles (Hagopian 2011, Lee 2014, Vatanen 2018, Stewart 2018), offer unique opportunities to explore microbial biomarker-disease relationships in well-characterized human populations. However, large cohort size and high sequencing cost make the full cohort biomarker study both expensive and inefficient. For instance, TEDDY has followed 8 676 newborns over 15 years to study the occurrence of persistent confirmed islet autoimmunity (IA, the pre-clinical phase of T1D) and the diagnosis of T1D, aiming to evaluate host genetics, gene expression, dietary biomarkers, metabolomics, microbiome, and virome in association to T1D. To efficiently study these biomarkers, TEDDY conducted the biomarker study in a nested case-control (NCC) sub-cohort (Lee 2014, Rundle 2012, Rundle 2015). Specifically, subjects with persistent IA and T1D are included as cases, while autoantibody-negative subjects at the case’s event age, matched by clinical center, sex, and family history of T1D are selected as controls to form the NCC sub-cohort. Only biospecimen from the NCC sub-cohort were processed to evaluate the corresponding biomarkers in relation to IA/T1D onset. The NCC design offers a cost-efficient solution for biomarker studies in a large cohort, but also introduces analytical complexities.

While the monthly to quarterly gut microbiome data are available in the TEDDY NCC sub-cohort, TEDDY study group has primarily analyzed microbiome data using cross-sectional approaches such as conditional logistic regression (CLR) (Vatanen 2018, Stewart 2018). These studies examined associations between microbial diversity or abundance at each sampling time points and dichotomous case–control status, and limited microbial associations with IA or T1D were detected. This highlights an urgent need for methods capable of jointly modeling longitudinally sampled, non-normally distributed microbial biomarker profiles and their temporal trajectories alongside the time to onset of IA/T1D, while accounting for the NCC design. Furthermore, The TEDDY study group reported the heterogeneity of T1D by categorizing IA into distinct phenotypes based on the first appearing autoantibody (IAA, GADA, IA2A) (Krischer 2015, Krischer 2017), underscoring the need for methods that account for competing risks in survival analysis to model the heterogeneity.

Several analytical methods have been developed for NCC studies, including the standard conditional logistic regression model for matched case-control data (Gail 1981) that the previous TEDDY microbiome association analyses have employed (Vatanen 2018, Stewart 2018), and refined methods such as the inverse selection probability weighting method (Samuelsen 1997), the local averaging method (Chen 2001), and likelihood-based methods (Scheike 2004, Zeng 2006). However, these methods cannot accommodate time-varying covariates such as repeatedly measured microbiome data generated by TEDDY. Joint modelling frameworks, which integrate longitudinal covariates and time-to-event outcomes, provide a promising alternative (Hogan 1997, Tsiatis 2004, Papageorgiou 2019). However, special considerations are required to extend such frameworks to NCC designs. Tseng and Liu (2009) first proposed a joint modeling approach for NCC studies, constructing a likelihood function based on all observed data to assess the relationships between longitudinal covariates and time-to-event outcomes. San (2013) extended this approach by focusing on NCC sub-cohort data only and employing an inverse probability weighting likelihood function for parameter estimation. Both methods are limited by their reliance on normality assumptions for longitudinal data and their inability to handle competing risks survival outcomes.

To our knowledge, no current method adequately addresses the following challenges: 1) characterizing longitudinally measured, non-normally distributed microbial biomarker data, 2) accommodating the NCC design while leveraging clinical metadata and survival outcome data from the full cohort, 3) modeling competing patterns in the appearance of autoantibodies.

Motivated by the unique TEDDY microbiome study, we propose a joint modeling method (JM-NCC) for analyzing longitudinal microbial biomarkers and competing events under NCC sampling. JM-NCC comprises two sub-models: a generalized linear mixed-effects model (longitudinal sub-model) to model biomarker trajectories over time, and a cause-specific hazard model (survival sub-model) to link these trajectories to competing events. We propose two inference approaches, i.e., fJM-NCC and wJM-NCC, depending on the availability of full cohort clinical metadata. The fJM-NCC approach integrates longitudinal measurements from the NCC sub-cohort with clinical metadata and survival outcome data from the full cohort to construct the likelihood function, thereby maximizing the use of available information. In contrast, wJM-NCC relies exclusively on NCC sub-cohort data and employs an inverse probability weighting likelihood function to address selection bias in NCC sampling. Additionally, a novel sandwich estimator is proposed in wJM-NCC to improve the estimation of parameter standard errors, addressing the underestimation issue in previous studies (San 2013). wJM-NCC serves as a useful alternative that complements fJM-NCC, particularly when access to full cohort clinical metadata is limited. **Figure 1** illustrates the NCC study design with TEDDY biomarker study as an example, and the data utilized by fJM-NCC and wJM-NCC.

**Figure 1:**
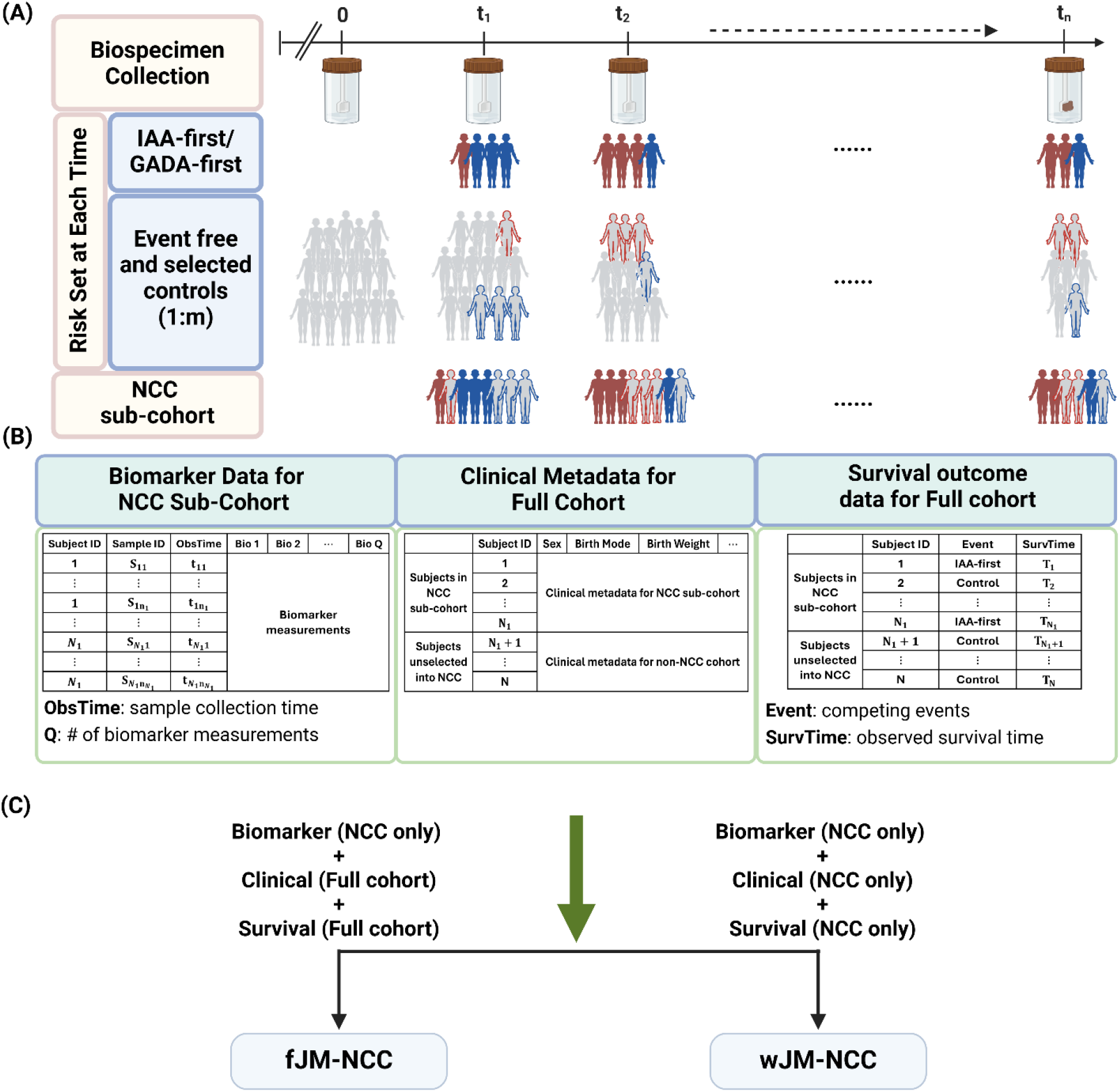
Overview of the Nested Case-Control (NCC) study design and data used by fJM-NCC and wJM-NCC, using TEDDY biomarker study as an example. **(A)** Full cohort biospecimen collection and construction of the NCC sub-cohort based on 1: *m* ratio. **(B)** Analyzed data consisting of three main components: longitudinal biomarker data available only in NCC sub-cohort, clinical metadata, and survival outcomes for the full cohort. **(C)** Data requirements for fJM-NCC and wJM-NCC. The fJM-NCC method requires NCC biomarker data, full cohort clinical data and survival outcomes while the wJM-NCC method only requires data from the NCC sub-cohort.

The remainder of this article is organized as follows: Section 2 introduces the notation and details of the joint modeling framework JM-NCC along with two inference approaches, fJM-NCC and wJM-NCC. Section 3 evaluates the parameter estimation and hypothesis testing performance of these two methods, comparing them with competing methods via the extensive simulation study. Section 4 illustrates the utility of fJM-NCC and wJM-NCC via longitudinal TEDDY microbiome study. Section 5 concludes with a discussion and future directions.

## 2 Methods

### 2.1 Notation and Model Specification

#### Full cohort

We consider a prospective cohort where the study outcomes are competing events, and longitudinal, high-dimensional microbial biomarkers are evaluated within a sub-cohort selected via nested case-control (NCC) sampling. Let the cohort consist of *N* subjects. Following the notation of Tseng and Liu (2009), let 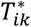 represent the time to competing event *k* (where *k* = 1, 2, …, *K*), and *C*_*i*_ denote the censoring time for subject *i*( where *i* = 1, 2, …, *N*). The observed survival time is defined by the observed survival time 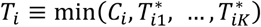, with the censoring indicator *δ*_*i*_ ∈ {0, 1, …, *K*}, where *δ*_*i*_ = 0 represents censoring, and *δ*_*i*_ = *k* specifies that subject *i* experienced competing event *k*. Baseline clinical metadata (covariates) for each subject *i* are denoted by ***X***_*i*_ = (*X*_*i*1_, *X*_*i*2_, …, *X*_*ip*_)^*T*^ of length *p*. The full cohort data are thus denoted as {(*T*_*i*_, *δ*_*i*_, ***X***_*i*_); *i* = 1, 2, …, *N*}.

#### NCC sub-cohort

All subjects with *δ*_*i*_ ≠ 0 are further included in the NCC sub-cohort as cases. For each case *i, m* (≥ 1 denotes the control-to-case ratio) controls are randomly selected from the corresponding risk set, consisting of individuals who have not experienced any event by time *T*_*i*_ and match case *i* on specific matching factors. Let the NCC sub-cohort consist of *N*_1_ (*N*_1_ < *N*) subjects. The statistical efficiency of the NCC study depends on the number of controls selected per case (*m*), with increasing numbers of controls approaching to the full-cohort statistical power. Longitudinal high-dimensional `-omics biomarker data are collected only for subjects in the NCC sub-cohort to retain study power while minimizing costs. These biomarkers are denoted as 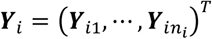 measured at times 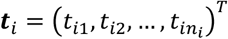 where 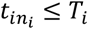. Using an indicator variable *R i* = 1 for subject *i* included in the NCC sub-cohort (*R*_*i*_ = 0 otherwise), the observed data for the NCC sub-cohort are denoted as {(*T*_*i*_, *δ*_*i*_, ***X***_*i*_, ***Y***_*i*_, ***t***_*i*_); *R*_*i*_ = 1}.

We propose a joint model framework **JM-NCC** for longitudinal and competing event outcomes under NCC sampling. JM-NCC comprises two sub-models: a generalized linear mixed-effects model for capturing the temporal trajectory of longitudinal biomarker profiles (the longitudinal sub-model) and a cause-specific hazard model (Lau 2009) for competing outcomes (the survival sub-model).

#### Longitudinal sub-model

We first model the change of longitudinal biomarker measurements over time using the generalized linear mixed-effects model to account for non-normally distributed biomarker data. Noticeably, the proposed framework considers each biomarker separately, and therefore we redefine 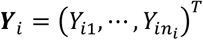 to denote the longitudinal measurement of a specific biomarker at time 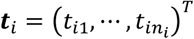 for simplicity. For a given biomarker, we assume the abundance of the *j*th sample *Y*_*ij*_ follows an exponential family distribution of *f*(*μ*_*ij*_, *a*_*ij*_(*τ*)*υ*(*μ*_*ij*_)) conditional on random effects *b*_*i*_ detailed below, where *μ*_*ij*_ = *E*[*Y*_*ij*_|*b*_*i*_] is the conditional mean of *Y*_*ij*_ given *b*_*i*_, *τ* is an unknown dispersion parameter, *a*_*ij*_(·) and *υ*( ·) are known functions (Dean 2007, Jiang 2007). Then we have

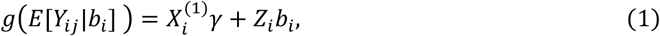

where *g*(·) is the link function relating the linear predictor 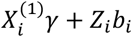 to the expectation *E*[*Y*_*ij*_|*b*_*i*_], 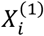 and *γ* correspond to the fixed effects design matrix that including times ***t***_*i*_ and a subset of baseline covariates, and coefficients, while *Z*_*i*_ and *b*_*i*_ denote the random effects design matrix and subject-specific random effects. Random effect *b*_*i*_ are assumed to follow a multivariate normal distribution *N*(0, Σ_*θ*_). Sub-model (1) simplifies to a linear mixed-effects model when *Y*_*ij*_ is assumed to follow a normal distribution *N*(*μ*_*ij*_, *σ*^2^), with parameters *a*_*ij*_(*τ*) = *σ*^2^ and *υ*(*μ*_*ij*_) = 1. For read count data with over-dispersion, we can model *Y*_*ij*_ using a negative binomial distribution *NB*(*μ*_*ij*_, *φ*) with mean *μ*_*ij*_ and dispersion parameters *φ*, where *a*_*ij*_(*τ*) = 1 and 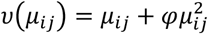.

#### Survival sub-model

The survival sub-model uses a cause-specific hazard model to estimate associations between longitudinal measurements and competing events:

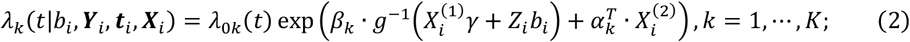

where *λ*_0*k*_(*t*) is the baseline cause-specific hazard function for competing event *k* (*k* = 1, 2, …, *K*), *g*^−1^(·) is the inverse of link function *g*(·), and 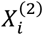 represents the subset of baseline covariates associated with the competing events. The regression coefficients *β*_*k*_ (*k* = 1, …, *K*) are the parameters of interest, representing the association between the temporal trajectory of the assessed biomarker and competing event *k*, while *α*_*k*_ represents the association between the covariates 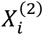 and competing event *k*. The longitudinal covariates in 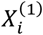 may overlap with survival covariates 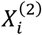, the interpretation of *α*_*k*_ can be considered as the effect of 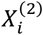 on competing event *k*, conditional on the true value of ***Y***, which is 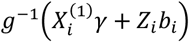. For *K* = 1, this model reduces to a Cox PH model for a general time-to-event outcome.

Let Φ = (*γ, τ*, vech(Σ_*θ*_), *λ*_0*k*_(*t*), *β*_*k*_, *α*_*k*_) be the set of all parameters to be estimated, where vech(·) denotes the half-vectorization of a matrix. The baseline hazard function *λ*_0*k*_(*t*) is modeled using a piecewise-constant approach: 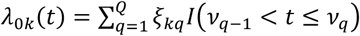, where 0 = *ν*_0_ < *ν*_1_ … < *ν*_*Q*_ denotes a split of the time scale and *ξ*_*kq*_ represents the hazard value for competing *k* within (*ν*_*q*−1_, *ν*_*q*_]. We split the parameter set Φ into two components, the parametric component *ϕ* = (*γ, τ*, vech(Σ_*θ*_), *β*_*k*_, *α*_*k*_) and the collection of non-parametric baseline hazard Λ_*k*_ = (*ξ*_*k*1_, …, *ξ*_*kQ*_). Our objective is to estimate both *ϕ* and Λ_*k*_. However, it is important to note that our primary interest lies in making inferences about *ϕ*.

In the following subsections, we present two MLE approaches for estimating *ϕ*, referred to as fJM-NCC and wJM-NCC respectively. The fJM-NCC method constructs the likelihood function by integrating longitudinal measurements from the NCC sub-cohort with survival data and clinical metadata from the full cohort. However, in many biomarker studies, certain clinical metadata, such as diet and blood concentrations of nutrient biomarkers in the TEDDY study, are not accessed for all subjects in the full cohort. To address this, the wJM-NCC method builds an inverse probability weighting likelihood function (Samuelsen 1997) using only the data from NCC sub-cohort for parameter estimation.

### 2.2 fJM-NCC: Likelihood Inference with Full Maximum Likelihood Approach

The fJM-NCC method incorporates all observed data and survival outcome to estimate *ϕ*, treating the unobserved longitudinal measurements as missing at random. The full log-likelihood function is given by

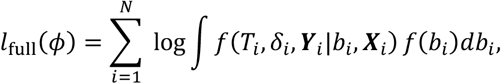

where *f*(*T*_*i*_, *δ*_*i*_, ***Y***_*i*_|*b*_*i*_, ***X***_*i*_) is the conditional probability density function (pdf) of (*T*_*i*_, *δ*_*i*_, ***Y***_*i*_) given *b*_*i*_ and ***X***_*i*_, *f*(*b*_*i*_) is the pdf of *b*_*i*_. Assuming the longitudinal and time-to-event data generating processes are conditionally independent given *b*_*i*_, the joint density simplifies to *f*(*T*_*i*_, *δ*_*i*_, ***Y***_*i*_|*b*_*i*_, ***X***_*i*_) = *f*(*T*_*i*_, *δ*_*i*_|*b*_*i*_, ***X***_*i*_)*f*(***Y***_*i*_|*b*_*i*_, ***X***_*i*_). For subjects with unobserved ***Y***_*i*_, {*i*; *R*_*i*_ = 0}, *f*(***Y***_*i*_|*b*_*i*_, ***X***_*i*_) = 1. Therefore, the full log-likelihood function can be rewritten as

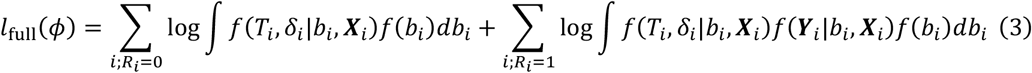

Based on the model specifications in Section 2.1, the pdfs of survival outcomes, random effects, and longitudinal data in the full log-likelihood function are expressed as follows:

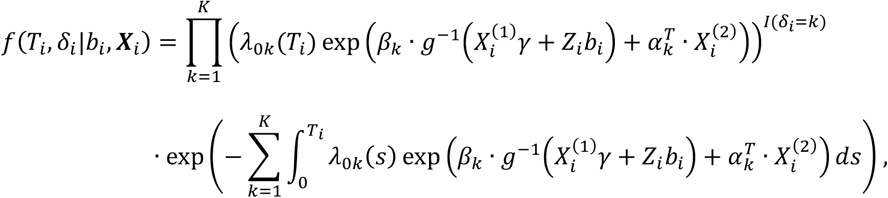

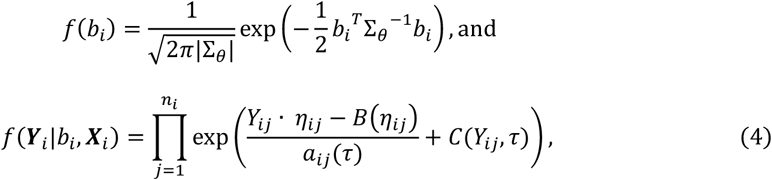

where *η*_*ij*_ (the nature parameter) is associated with the conditional mean *μ*_*ij*_, *B*(·) and C(·) are known distribution-specific functions (Jiang 2007). Gaussian quadrature is employed to approximate the integral in the function (3). The MLEs of *ϕ* are obtained by maximizing the full log-likelihood function (3). The standard errors (SE) of parameters are derived using the Fisher’s information matrix, which is consistently estimated by the empirical Fisher’s information matrix. The 95% confidence intervals (CIs) of the parameters are calculated as 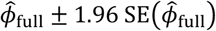, where SE(·) is based on the Fisher’s information matrix. The Wald test statistic (Gregory 1985) is used to test this null hypothesis for each competing event *k*: *H*_0_: *β*_*k*_ = 0.

### 2.3 wJM-NCC: Likelihood Inference with the Inverse Probability Weighting Approach

The wJM-NCC method constructs an inverse probability weighting likelihood function using only data from the NCC sub-cohort, accounting for the potential selection bias in the NCC sampling. Such bias arises because subjects who remain in the full cohort for longer duration are more likely to be included in the NCC sub-cohort. The weighted log-likelihood function is given by:

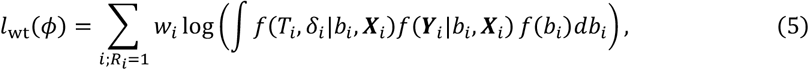

where weight 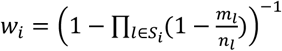 presents the inverse probability of inclusion as a control. *S*_*i*_ is the set of cases for which subject *i* was eligible to be selected as a control, *m*_*l*_ is the number of controls selected for case *l*, and *n*_*l*_ is the number of candidates in the risk set for case *l*. Notably, *w*_*i*_ = 1 if subject *i* is a case.

As in the full log-likelihood function, the integral component in function (5) is approximated using Gaussian quadrature to obtain the MLE of parameter *ϕ*.

#### Standard Error Estimation

Direct application of the Fisher’s information matrix to the weighted log-likelihood function tends to underestimate the standard errors (San 2013). To address this, we employed the sandwich method to derive a robust covariance estimator for the weighted log-likelihood function (5). The sandwich covariance estimator is given by

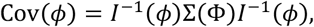

where

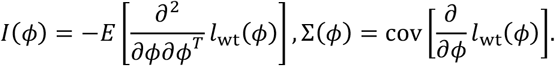

*I*(*ϕ*) and Σ(*ϕ*) can be consistently estimated by the empirical Fisher’s information matrix and the empirical covariance matrix, respectively:

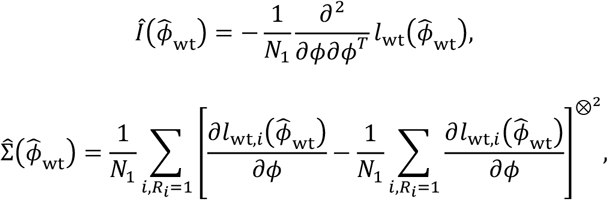

where 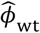 is the MLE of *ϕ* obtained by the wJM-NCC approach, and 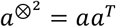 for any vector *a*. The 95% CIs for the parameters and hypothesis tests are conducted in the same manner as for fJM-NCC.

## 3 Simulation Studies

We evaluated the inference performance of the two proposed methods **fJM-NCC** and **wJM-NCC** respectively for joint modeling of longitudinal and competing event outcomes under the NCC design and compared these methods with alternative approaches using synthetic data.

### 3.1 Simulation Setup

- **Generating synthetic data for the full cohort:** We first simulated longitudinal biomarker measurements using a linear mixed-effects model with a fixed slope and random intercept: *Y*_*ij*_ = *γ* · *t*_*ij*_+ *b*_0*i*_ + *ϵ*_*ij*_. Each subject had five repeated measurements (*j* = 0,1, …, 4) recorded at times *t*_*ij*_=0,0.1,0.2,0.3,0.4 with fixed slope *γ* = 0.1. The random intercept *b*_0*i*_ was drawn from a normal distribution *N*(0, θ^2^ = 2), and the error term *ϵ*_*ij*_~*N*(0, *σ*^2^ = 1). In real-world scenarios, longitudinal biomarker data may not be available for all subjects. The next sub-section describes how we sample a synthetic NCC sub-cohort dataset from the full cohort. To simulate competing event times, we considered a dichotomous covariate, gender 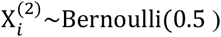, with a coefficient *α* = −0.2 in the cause-specific hazard model for competing risks, defined as 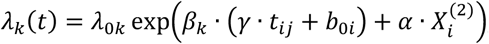, where we considered two competing events ( *k* = 1, 2) with constant baseline hazards *λ*_01_ = *e*^−5^ and *λ*_02_ = *e*^−4^ respectively. The parameter *β*_*k*_ (*k* = 1,2) represents the association between longitudinal biomarker trajectories and the competing event outcomes, which is the central focus of our study.
- **Simulation Scenarios:** We considered two scenarios to evaluate the performance of the proposed methods in the statistical inference of *β*_1_ and *β*_2_:
  ▪ **Scenario 1:** Global Null Hypothesis: *β*_1_ = *β*_2_ = 0, indicating no association between the biomarker trajectory and either event.
  ▪ **Scenario 2:** Alternative Hypothesis: In this scenario, we fixed *β*_2_ = 0.1 and varied *β*_1_ from 0 (no association with event 1, no competing) to 0.3, with an increment of 0.01. This allows us to examine performance under alternative hypotheses with varying effect sizes for event 1.
- **Sampling an NCC sub-cohort:** To replicate the event rate of islet autoimmunity (IA) observed in the TEDDY study (Tsiatis 2004), we simulated competing events data for *N* = 8,000 individuals. The first 400 individuals were selected based on the earliest event time for either event 1 or event 2, and form the cases in the NCC sub-cohort. The maximum event time observed among these 400 individuals was then used as the censoring time for the remaining participants. For each case, we selected *m* = 1, 3, or 5 controls from the corresponding risk set, matched by gender 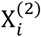, to construct the NCC sub-cohort. Covariates, longitudinal and survival outcome data were then extracted for each selected individual to form the NCC dataset, resulting in sub-cohort sizes of 10% (m=1), 20% (m=3), and 30% (m=5) of the full cohort size.

### 3.2 Competing Methods

- **Oracle method:** In the simulation studies, covariates, longitudinal, and survival outcomes data were generated for all individuals in the full cohort. As a benchmark, we used a joint modeling approach with competing events modeled that assumes the full cohort data, i.e., covariates, longitudinal and survival outcomes are available for all subjects. This approach, denoted as the **Oracle** method, uses the log-likelihood function 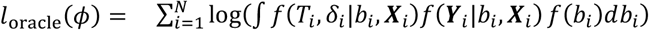 for parameter estimation and hypothesis testing, where the pdfs are defined in Equation (4). The Oracle method serves as a baseline to highlight the benefits of our methods designed for NCC sampling. While the Oracle method relies on the full cohort data, our NCC-based methods reduce sample size requirements for biomarker studies and achieve minimal efficiency loss compared to the Oracle method in parameter estimation and hypothesis testing.
- **wJM-NCC(Fisher) method:** To evaluate standard error estimation calibration in our proposed wJM-NCC method, we also included the weighted likelihood approach used in wJM-NCC, but with standard error estimates derived from the Fisher’s information matrix. We denote this approach as **wJM-NCC(Fisher)**. Note that the point estimates from both wJM-NCC and wJM-NCC(Fisher) are the same; the only difference lies in the standard error estimates for the studied parameters. This comparison highlights the underestimation of standard errors in the Fisher method, which the sandwich estimator in wJM-NCC avoids.
- **JM method:** Additionally, we included the classical joint modeling approach for longitudinal outcomes and competing events, as proposed by Williamson (2008). This approach assumes a Gaussian linear model for longitudinal outcomes, and a semi-parametric cause-specific hazard model for the competing survival outcomes. Implemented in the R package JM (Rizopoulos 2010), the method can be executed using the function jointModel(lmeObject, survObject, timeVar=“obsTime”, method=“spline-PH-GH”, CompRisk = TRUE, interFact = list(value = ~ causeEvent, data = cc.data.long)), where cc.data.long is the competing risks long format data of NCC sub-cohort obtained using function crLong(). The matching variable 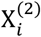 is included as a covariate, similar to Oracle, fJM-NCC, wJM-NCC, and wJM-NCC(Fisher). We refer to this method as **JM** in this article. Since JM requires both longitudinal and survival data for each individual included in the model, it can only be applied to the NCC sub-cohort, as longitudinal measurements are unavailable for individuals not selected into the NCC sub-cohort (i.e., where *R*_*i*_ = 0).
- **CLR method:** Conditional logistic regression (**CLR**) is a standard approach for analyzing matched case-control data to study the relationship between risk factors and a dichotomous outcome with strata (Gail 1981). Given that NCC sampling with matching can be considered a matched case-control design -- and has been applied to analyze TEDDY data (Norris 2018, Lee 2020, Auchtung 2022) -- CLR was included in our comparison with some necessary adaptations. First, because CLR cannot accommodate time-varying covariates (e.g., longitudinal biomarker measurements), we included each individual’s average biomarker measurement as a covariate. Second, cases and their matched controls were analyzed for each type of competing event to estimate parameters specific to that event. Thus, CLR results reflect the association between average biomarker values and each individual event, rather than a direct inference on competing risks.

**Table 1** summarizes all methods considered in this article, outlining their data requirements, applicable study designs, and other relevant characteristics. Bias and mean squared error (MSE) of point estimates, and the corresponding mean standard errors (SE) and empirical standard errors (ESE) are employed to evaluate the point estimates of model parameters. Confidence Intervals (CI) estimates were assessed using the average length of 95% confidence intervals (CI-L) and empirical coverage probability (ECP). We further evaluated Type-I error rates and statistical power for hypothesis tests. A total of 1000 repetitions were conducted to evaluate the parameter estimates and hypothesis testing performance for each parameter combination, with a nominal significance level of 0.05.

**Table 1:**
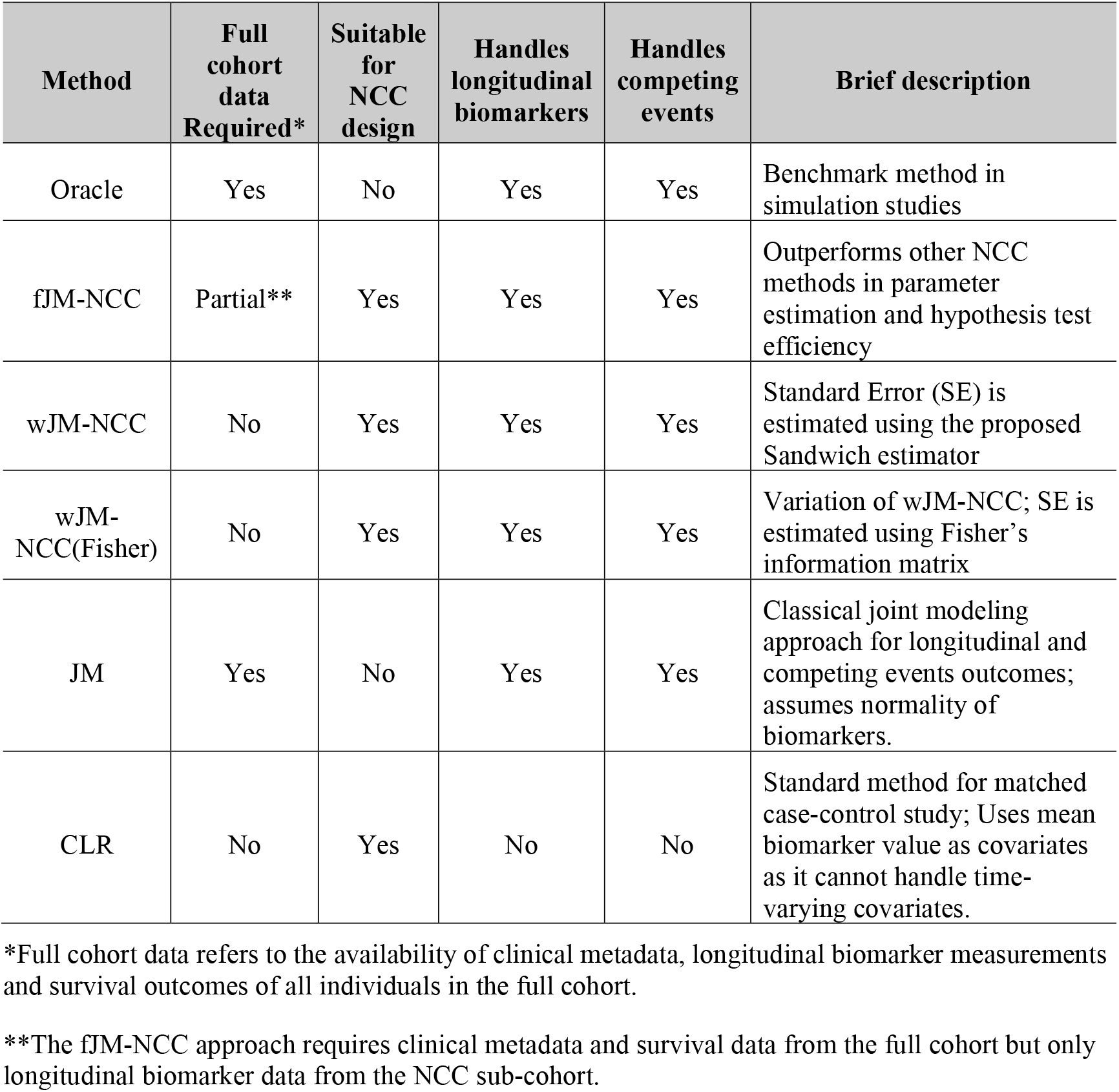
Summary of data requirements and applicability of methods to NCC design, longitudinal biomarkers, and competing events.

### 3.3 Simulation Results

**Table 2** summarizes the performance of point estimates and 95% confidence interval (CI) estimates for *β*_1_ and *β*_2_ under the global null hypothesis (*β*_1_ = *β*_2_ = 0, **Scenario 1**). Across all methods, the point estimates for *β*_1_ and *β*_2_ are nearly unbiased. The standard error (SE) estimates for both parameters are consistent for all methods except wJM-NCC(Fisher), as evidenced by their average SEs closely matching the empirical standard errors (ESEs), while the SEs from wJM-NCC(Fisher) are smaller than the corresponding ESEs. The newly proposed sandwich estimator in wJM-NCC provides well-calibrated SE estimates.

**Table 2:** Performance of all methods for point and 95% confidence interval estimation of *β*_1_ and *β*_2_ under **Scenario 1** ( *β*_1_ = *β*_2_ = 0).

| $m^1$ | Method | $\beta_1$ | | | | | | $\beta_2$ | | | | | |
| --- | --- | --- | --- | --- | --- | --- | --- | --- | --- | --- | --- | --- | --- |
|  |  | Bias | SE<sup>2</sup> | ESE<sup>3</sup> | MSE<sup>4</sup> | CI-L<sup>5</sup> | ECP<sup>6</sup> | Bias | SE | ESE | MSE | CI-L | ECP |
| 1 | Oracle | -0.002 | 0.073 | 0.071 | 0.005 | 0.285 | 0.958 | -0.001 | 0.044 | 0.046 | 0.002 | 0.172 | 0.929 |
|  | fJM-NCC | -0.003 | 0.074 | 0.073 | 0.005 | 0.292 | 0.959 | -0.001 | 0.046 | 0.049 | 0.002 | 0.180 | 0.935 |
|  | wJM-NCC | -0.003 | 0.082 | 0.081 | 0.007 | 0.320 | 0.950 | -0.002 | 0.057 | 0.060 | 0.004 | 0.225 | 0.939 |
|  | wJM-NCC(Fisher) | -0.003 | 0.073 | 0.081 | 0.007 | 0.285 | 0.920 | -0.002 | 0.044 | 0.060 | 0.004 | 0.172 | 0.857 |
|  | JM | -0.003 | 0.072 | 0.072 | 0.005 | 0.284 | 0.958 | -0.002 | 0.044 | 0.046 | 0.002 | 0.171 | 0.937 |
|  | CLR | -0.003 | 0.097 | 0.095 | 0.009 | 0.380 | 0.954 | -0.004 | 0.058 | 0.059 | 0.004 | 0.226 | 0.942 |
| 3 | Oracle | -0.001 | 0.072 | 0.072 | 0.005 | 0.284 | 0.954 | 0.001 | 0.044 | 0.043 | 0.002 | 0.172 | 0.957 |
|  | fJM-NCC | 0.000 | 0.073 | 0.073 | 0.005 | 0.288 | 0.955 | 0.001 | 0.045 | 0.044 | 0.002 | 0.177 | 0.958 |
|  | wJM-NCC | 0.000 | 0.075 | 0.075 | 0.006 | 0.293 | 0.961 | 0.001 | 0.048 | 0.047 | 0.002 | 0.190 | 0.960 |
|  | wJM-NCC(Fisher) | 0.000 | 0.072 | 0.075 | 0.006 | 0.284 | 0.946 | 0.001 | 0.044 | 0.047 | 0.002 | 0.172 | 0.938 |
|  | JM | -0.001 | 0.072 | 0.071 | 0.005 | 0.282 | 0.955 | 0.001 | 0.044 | 0.043 | 0.002 | 0.171 | 0.960 |
|  | CLR | 0.000 | 0.077 | 0.080 | 0.006 | 0.303 | 0.948 | -0.001 | 0.047 | 0.046 | 0.002 | 0.183 | 0.957 |
| 5 | Oracle | 0.003 | 0.072 | 0.072 | 0.005 | 0.284 | 0.961 | 0.000 | 0.044 | 0.044 | 0.002 | 0.172 | 0.950 |
|  | fJM-NCC | 0.002 | 0.073 | 0.074 | 0.005 | 0.287 | 0.957 | 0.000 | 0.045 | 0.045 | 0.002 | 0.176 | 0.960 |
|  | wJM-NCC | 0.002 | 0.074 | 0.074 | 0.006 | 0.289 | 0.959 | 0.000 | 0.046 | 0.046 | 0.002 | 0.182 | 0.957 |
|  | wJM-NCC(Fisher) | 0.002 | 0.072 | 0.074 | 0.006 | 0.284 | 0.951 | 0.000 | 0.044 | 0.046 | 0.002 | 0.172 | 0.942 |
|  | JM | 0.002 | 0.072 | 0.072 | 0.005 | 0.282 | 0.956 | -0.001 | 0.044 | 0.044 | 0.002 | 0.171 | 0.955 |
|  | CLR | 0.000 | 0.073 | 0.073 | 0.005 | 0.287 | 0.953 | -0.001 | 0.044 | 0.044 | 0.002 | 0.174 | 0.946 |

1. Control-to-case ratio, i.e., the number of controls per case in the NCC sub-cohort
2. Estimated standard error
3. Empirical standard error
4. Mean squared error
5. Average length of the 95% confidence intervals
6. Empirical coverage probability of the 95% confidence interval

For 95% CI estimation, evaluated by average CI length and empirical coverage probability (ECP), fJM-NCC, wJM-NCC, and JM yield comparable results to Oracle across varying values of *m*. However, wJM-NCC(Fisher) produces shorter CIs due to SE underestimation, leading to lower coverage rates (ECP). The CLR method generates the longest CIs across most settings.

**Tables 3-4** and **Tables S1-S2** present the results for point and 95% CI estimates for *β*_1_ and *β*_2_, with *β*_2_ fixed at 0.1 and *β*_1_ varying from 0 (**Table 3**), 0.1 (**Table S1**), 0.2 (**Table S2**), to 0.3 (**Table 4**) under **Scenario 2**. The point estimates for *β*_1_ and *β*_2_ from Oracle, fJM-NCC, and wJM-NCC remain unbiased with minimal bias, while JM shows considerable bias across all parameter settings. This bias arises because JM incorrectly treats the NCC sub-cohort as the full cohort, despite the event rate in the NCC sub-cohort being distorted to 1/(*m* + 1). The bias decreases as the number of controls per case (*m*) increases, making the event rate in the NCC sub-cohort more representative of the full cohort. The CLR method provides unbiased point estimates for *β*_1_, but exhibits substantial bias for *β*_2_ when *β*_1_ = 0 and *β*_2_ = 0.1 (Table 3), where the biomarker is only associated with one event. The bias in CLR’s point estimate for *β*_1_ becomes more pronounced when biomarker value is associated with competing events (both *β*_1_ and *β*_2_ are non-zero; Tables 4, S1-S2). SE estimates for *β*_1_ and *β*_2_ are consistent across Oracle, fJM-NCC, wJM-NCC, closely aligning with the ESEs. However, wJM-NCC (Fisher) continues to underestimate SEs. The 95% CI estimation performances of fJM-NCC, wJM-NCC, wJM-NCC(Fisher), and CLR in Scenario 2 are similar to those observed in Scenario 1.

**Table 3:** Performance of all methods for point and 95% confidence interval estimation of *β*_1_ and *β*_2_ under **Scenario 2** (*β*_1_ = 0 and *β*_2_ = 0.1).

| $m^1$ | Method | $\beta_1$ | | | | | | $\beta_2$ | | | | | |
| --- | --- | --- | --- | --- | --- | --- | --- | --- | --- | --- | --- | --- | --- |
|  |  | Bias | SE<sup>2</sup> | ESE<sup>3</sup> | MSE<sup>4</sup> | CI-L<sup>5</sup> | ECP<sup>6</sup> | Bias | SE | ESE | MSE | CI-L | ECP |
| 1 | Oracle | 0.001 | 0.073 | 0.074 | 0.006 | 0.287 | 0.944 | 0.000 | 0.044 | 0.043 | 0.002 | 0.171 | 0.946 |
|  | fJM-NCC | 0.001 | 0.075 | 0.077 | 0.006 | 0.294 | 0.950 | 0.002 | 0.046 | 0.046 | 0.002 | 0.182 | 0.946 |
|  | wJM-NCC | 0.003 | 0.082 | 0.083 | 0.007 | 0.322 | 0.950 | 0.003 | 0.058 | 0.059 | 0.004 | 0.227 | 0.946 |
|  | wJM-NCC(Fisher) | 0.003 | 0.073 | 0.083 | 0.007 | 0.288 | 0.929 | 0.003 | 0.044 | 0.059 | 0.004 | 0.172 | 0.850 |
|  | JM | -0.019 | 0.073 | 0.073 | 0.006 | 0.285 | 0.938 | -0.022 | 0.044 | 0.045 | 0.002 | 0.171 | 0.914 |
|  | CLR | 0.003 | 0.098 | 0.101 | 0.010 | 0.384 | 0.956 | -0.009 | 0.058 | 0.058 | 0.003 | 0.228 | 0.949 |
| 3 | Oracle | 0.002 | 0.073 | 0.075 | 0.006 | 0.287 | 0.945 | 0.001 | 0.044 | 0.044 | 0.002 | 0.171 | 0.946 |
|  | fJM-NCC | 0.002 | 0.074 | 0.078 | 0.006 | 0.291 | 0.945 | 0.002 | 0.045 | 0.046 | 0.002 | 0.178 | 0.957 |
|  | wJM-NCC | 0.001 | 0.076 | 0.079 | 0.006 | 0.298 | 0.943 | 0.000 | 0.049 | 0.050 | 0.002 | 0.190 | 0.946 |
|  | wJM-NCC(Fisher) | 0.001 | 0.073 | 0.079 | 0.006 | 0.287 | 0.937 | 0.000 | 0.044 | 0.050 | 0.002 | 0.172 | 0.921 |
|  | JM | -0.008 | 0.073 | 0.075 | 0.006 | 0.285 | 0.944 | -0.011 | 0.044 | 0.044 | 0.002 | 0.171 | 0.941 |
|  | CLR | 0.004 | 0.078 | 0.081 | 0.007 | 0.306 | 0.935 | -0.008 | 0.047 | 0.048 | 0.002 | 0.184 | 0.945 |
| 5 | Oracle | -0.005 | 0.073 | 0.074 | 0.005 | 0.286 | 0.940 | 0.001 | 0.044 | 0.043 | 0.002 | 0.171 | 0.953 |
|  | fJM-NCC | -0.005 | 0.074 | 0.075 | 0.006 | 0.290 | 0.939 | 0.002 | 0.045 | 0.045 | 0.002 | 0.176 | 0.955 |
|  | wJM-NCC | -0.005 | 0.074 | 0.076 | 0.006 | 0.291 | 0.941 | 0.001 | 0.046 | 0.046 | 0.002 | 0.182 | 0.945 |
|  | wJM-NCC(Fisher) | -0.005 | 0.073 | 0.076 | 0.006 | 0.286 | 0.936 | 0.001 | 0.044 | 0.046 | 0.002 | 0.172 | 0.927 |
|  | JM | -0.010 | 0.072 | 0.074 | 0.006 | 0.284 | 0.938 | -0.007 | 0.043 | 0.043 | 0.002 | 0.170 | 0.947 |
|  | CLR | -0.003 | 0.074 | 0.075 | 0.006 | 0.289 | 0.942 | -0.008 | 0.044 | 0.045 | 0.002 | 0.174 | 0.945 |

1. Control-to-case ratio, i.e., the number of controls per case in the NCC sub-cohort
2. Estimated standard error
3. Empirical standard error
4. Mean squared error
5. Average length of the 95% confidence intervals
6. Empirical coverage probability of the 95% confidence interval

**Table 4:** Performance of all methods for point and 95% confidence interval estimation of *β*_1_ and *β*_2_ under **Scenario 2** (*β*_1_ = 0.3 and *β*_2_ = 0.1).

| $m^1$ | Method | $\beta_1$ | | | | | | $\beta_2$ | | | | | |
| --- | --- | --- | --- | --- | --- | --- | --- | --- | --- | --- | --- | --- | --- |
|  |  | Bias | SE<sup>2</sup> | ESE<sup>3</sup> | MSE<sup>4</sup> | CI-L<sup>5</sup> | ECP<sup>6</sup> | Bias | SE | ESE | MSE | CI-L | ECP |
| 1 | Oracle | -0.001 | 0.070 | 0.074 | 0.005 | 0.275 | 0.938 | 0.001 | 0.044 | 0.045 | 0.002 | 0.174 | 0.948 |
|  | fJM-NCC | 0.003 | 0.074 | 0.078 | 0.006 | 0.289 | 0.943 | 0.002 | 0.047 | 0.048 | 0.002 | 0.184 | 0.951 |
|  | wJM-NCC | 0.000 | 0.083 | 0.086 | 0.007 | 0.326 | 0.946 | 0.000 | 0.058 | 0.059 | 0.003 | 0.228 | 0.948 |
|  | wJM-NCC(Fisher) | 0.000 | 0.070 | 0.086 | 0.007 | 0.276 | 0.904 | 0.000 | 0.045 | 0.059 | 0.003 | 0.175 | 0.865 |
|  | JM | -0.054 | 0.070 | 0.071 | 0.008 | 0.273 | 0.872 | -0.048 | 0.044 | 0.045 | 0.004 | 0.172 | 0.797 |
|  | CLR | -0.022 | 0.101 | 0.103 | 0.011 | 0.396 | 0.936 | -0.005 | 0.059 | 0.061 | 0.004 | 0.232 | 0.949 |
| 3 | Oracle | 0.004 | 0.070 | 0.070 | 0.005 | 0.274 | 0.957 | 0.003 | 0.045 | 0.044 | 0.002 | 0.175 | 0.955 |
|  | fJM-NCC | 0.007 | 0.072 | 0.073 | 0.005 | 0.282 | 0.961 | 0.004 | 0.046 | 0.045 | 0.002 | 0.180 | 0.947 |
|  | wJM-NCC | 0.005 | 0.074 | 0.074 | 0.006 | 0.290 | 0.955 | 0.002 | 0.049 | 0.048 | 0.002 | 0.193 | 0.955 |
|  | wJM-NCC(Fisher) | 0.005 | 0.070 | 0.074 | 0.006 | 0.274 | 0.948 | 0.002 | 0.045 | 0.048 | 0.002 | 0.175 | 0.925 |
|  | JM | -0.023 | 0.069 | 0.069 | 0.005 | 0.272 | 0.945 | -0.020 | 0.044 | 0.043 | 0.002 | 0.173 | 0.929 |
|  | CLR | -0.021 | 0.078 | 0.079 | 0.007 | 0.306 | 0.938 | -0.005 | 0.048 | 0.047 | 0.002 | 0.187 | 0.949 |
| 5 | Oracle | 0.001 | 0.070 | 0.071 | 0.005 | 0.274 | 0.954 | 0.002 | 0.045 | 0.046 | 0.002 | 0.175 | 0.941 |
|  | fJM-NCC | 0.003 | 0.071 | 0.072 | 0.005 | 0.280 | 0.950 | 0.002 | 0.046 | 0.048 | 0.002 | 0.179 | 0.939 |
|  | wJM-NCC | 0.002 | 0.072 | 0.073 | 0.005 | 0.284 | 0.947 | 0.002 | 0.047 | 0.049 | 0.002 | 0.185 | 0.940 |
|  | wJM-NCC(Fisher) | 0.002 | 0.070 | 0.073 | 0.005 | 0.274 | 0.947 | 0.002 | 0.045 | 0.049 | 0.002 | 0.175 | 0.922 |
|  | JM | -0.018 | 0.069 | 0.070 | 0.005 | 0.272 | 0.947 | -0.013 | 0.044 | 0.046 | 0.002 | 0.173 | 0.934 |
|  | CLR | -0.023 | 0.073 | 0.073 | 0.006 | 0.286 | 0.937 | -0.006 | 0.045 | 0.047 | 0.002 | 0.177 | 0.939 |

1. Control-to-case ratio, i.e., the number of controls per case in the NCC sub-cohort
2. Estimated standard error
3. Empirical standard error
4. Mean squared error
5. Average length of the 95% confidence intervals
6. Empirical coverage probability of the 95% confidence interval

**Tables S3-S7** presents estimates for other parameters, including the fixed slope *γ*, the standard deviation (log) of random intercept log(*θ*), the standard deviation (log) of random error log(σ) in the longitudinal sub-model, and the fixed effect *α* of the covariate in the survival sub-model. CLR is excluded as it does not provide estimates for these parameters. JM does not provide SE estimates for log(*θ*) and log (*σ*) and therefore performance of JM for these parameters are not reported. Oracle, fJM-NCC, and wJM-NCC, as expected, yield minimal biased point estimates for all parameters. In contrast, JM shows bias for log(*θ*), log (*σ*), and *α* due to that it treats NCC sub-cohort as the full cohort. SE estimates from Oracle and fJM-NCC consistently align closely with their corresponding ESEs, while wJM-NCC provides consistent SE estimates for the longitudinal data generation parameters ( *γ*, log(*θ*), and log(σ)) but slightly conservative SEs for *α*. wJM-NCC(Fisher) underestimate SEs for for *γ*, log(*θ*), and log(σ). JM exhibits a consistent SE estimate for *γ* but a significant conservative SE estimate for *α*. In summary, these results indicate that JM and CLR are not suitable for analyzing NCC designed longitudinal biomarker studies with competing events.

**Figure 2 (A)** shows the empirical Type-I error rates for testing *H*_0_: *β*_1_ = 0 under **Scenario 1**. Results for testing *H*_0_: *β*_2_ = 0 are similar and omitted for brevity. Oracle, fJM-NCC, and wJM-NCC maintain Type-I error rates close to the nominal level of 0.05, demonstrating the validity of the proposed methods. wJM-NCC(Fisher) demonstrates inflated Type-I error rate due to SE underestimation, and its statistical power is excluded from further comparisons. JM shows elevated Type-I error rates when *m* = 1, where the event rate in the fitted NCC sub-cohort is most severely distorted. CLR also shows a higher-than-nominal Type-I error rate when *m* = 3.

**Figure 2:**
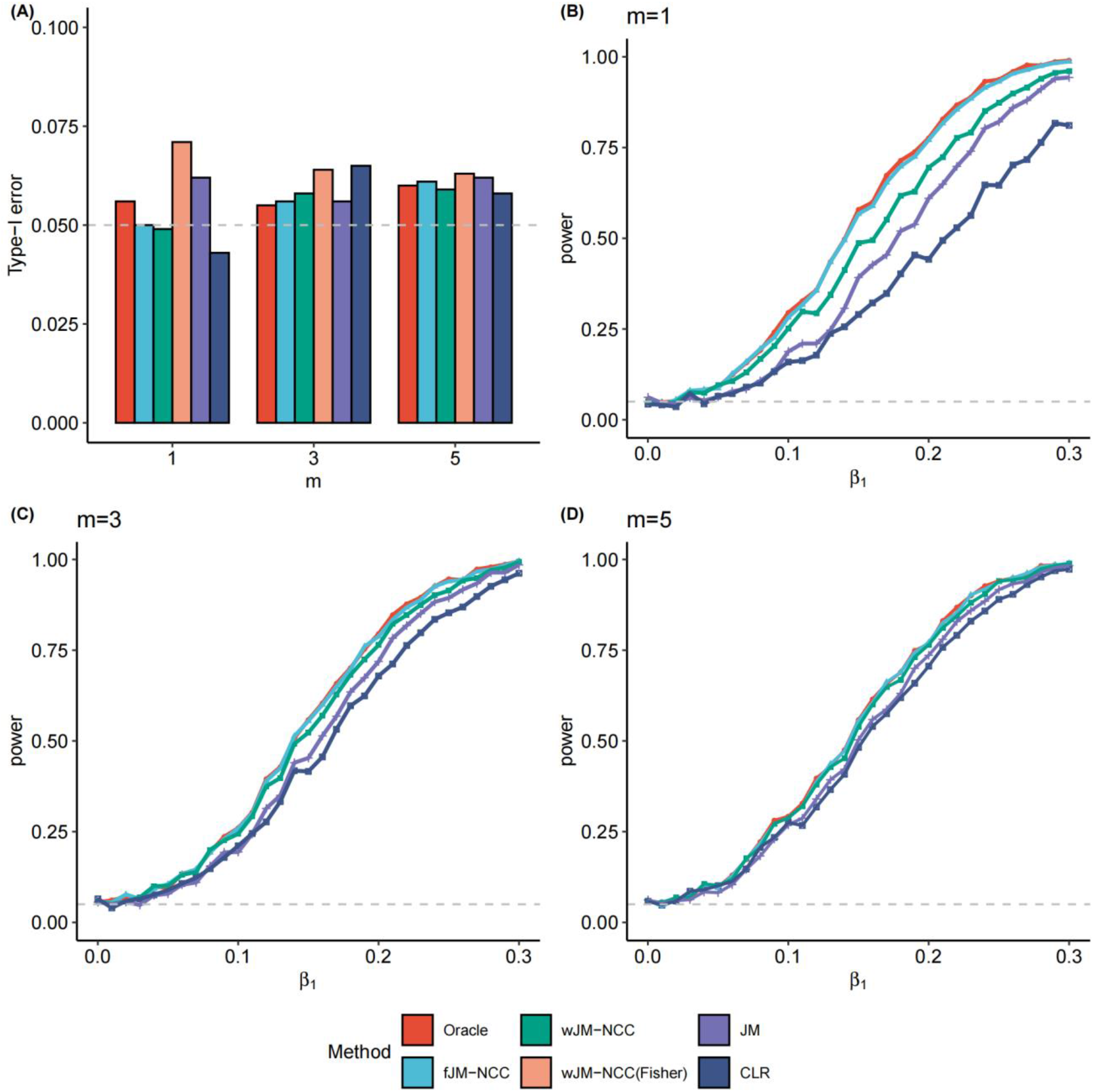
Statistical testing performance of all considered methods for testing *H*_0_: *β*_1_ = 0 under two scenarios. (**A)** Scenario 1: Empirical Type-I error rates of all methods for testing *H*_0_: *β*_1_ = 0 with *m* (control-to-case ratio) ranging from 1 to 5. (**B)-(D)** Scenario 2: Statistical power of all methods (excluding wJM-NCC(Fisher) due to inflated Type-I error) for *β*_1_ values ranging from 0 to 0.3 in increment of 0.01, with *m* = 1, 3, and 5, and *β*_2_ fixed at 0.1.

**Figure 2(B)-(D)** illustrates the statistical power for testing *H*_0_: *β*_1_ = 0 as *β*_1_ increases from 0 to 0.3 with increments of 0.01, and *β*_2_ fixed at 0.1 and *m* varying from 1 to 5, representing increasing sample sizes of the NCC sub-cohort (**Scenario 2**). Oracle, modeled on the full dataset, serves as the benchmark method and remains its results unchanged as *m* varies. As expected, Oracle achieves the highest statistical power under all scenarios. fJM-NCC closely follows Oracle, with nearly equivalent power across various values of *β*_1_. The statistical power of wJM-NCC ranks third, but still yields satisfactory results, particularly when compared to the traditional methods JM and CLR. The power loss due to NCC sampling becomes negligible when *m* = 5, as evidenced by nearly identical statistical powers for Oracle, fJM-NCC, and wJM-NCC.

These results indicate that fJM-NCC performs optimally when longitudinal biomarker data for the NCC sub-cohort and full cohort clinical metadata and survival outcomes are available. Nevertheless, wJM-NCC offers a complementary approach and achieves satisfactory performance, especially when full cohort clinical metadata is not available or difficult to access. Meanwhile, both JM and CLR consistently demonstrate lower statistical power compared to both Oracle method and the proposed methods fJM-NCC and wJM-NCC.

## 4 Application to TEDDY Microbiome Study

Here, we apply the proposed methods, fJM-NCC and wJM-NCC, and competing methods JM and CLR, to investigate the longitudinal microbiome profiles during early human life and their association with the competing appearance of two dominant autoantibodies, IAA (IAA-first) and GADA (GADA-first), using data from the comprehensive TEDDY microbiome biomarker study (Stewart 2018). Due to its inflated type I error rate in hypothesis testing, wJM-NCC(Fisher) was excluded from the analysis.

For this secondary data analysis, clinical metadata accessible to our research group were extracted from the full TEDDY cohort (N = 8,607), including sex, birth mode, birth weight, and the status of having a first-degree relative with T1D (FDR). Missing data were imputed using a binormal distribution for birth mode and FDR, and a normal distribution for birth weight, with parameters estimated from subjects with complete data. Microbiome data included community-level measurements and temporal shotgun metagenomic species-level taxa abundances from the NCC-sub cohort of 357 children who developed IAA first (N= 244) or GADA first (N = 113), along with their matched controls (N = 327). Community-level microbiome measurements include four alpha diversity indices -- microbial richness (number of OTUs/species) and Shannon’s diversity indices calculated from both 16S rRNA sequencing and shotgun metagenomic sequencing data -- and two microbiome maturation indices: Microbiota-by-age Z-scores (MAZ) and microbiota age derived from 16S rRNA sequencing data (Stewart 2018, Subramanian 2014). Data filtering steps are performed at both the microbiome sample and species level for the shotgun metagenomic taxa abundances. Microbiome samples with >10% missingness or zero abundances across all species were excluded, and species detected in fewer than 10% of samples or with an average relative abundance <0.01% were removed (Figure 3A). After these quality control steps, the relative abundances (arcsine square root transformed) of 231 species from 11,021 samples (819 subjects) are retained for downstream association analysis.

**Figure 3.**
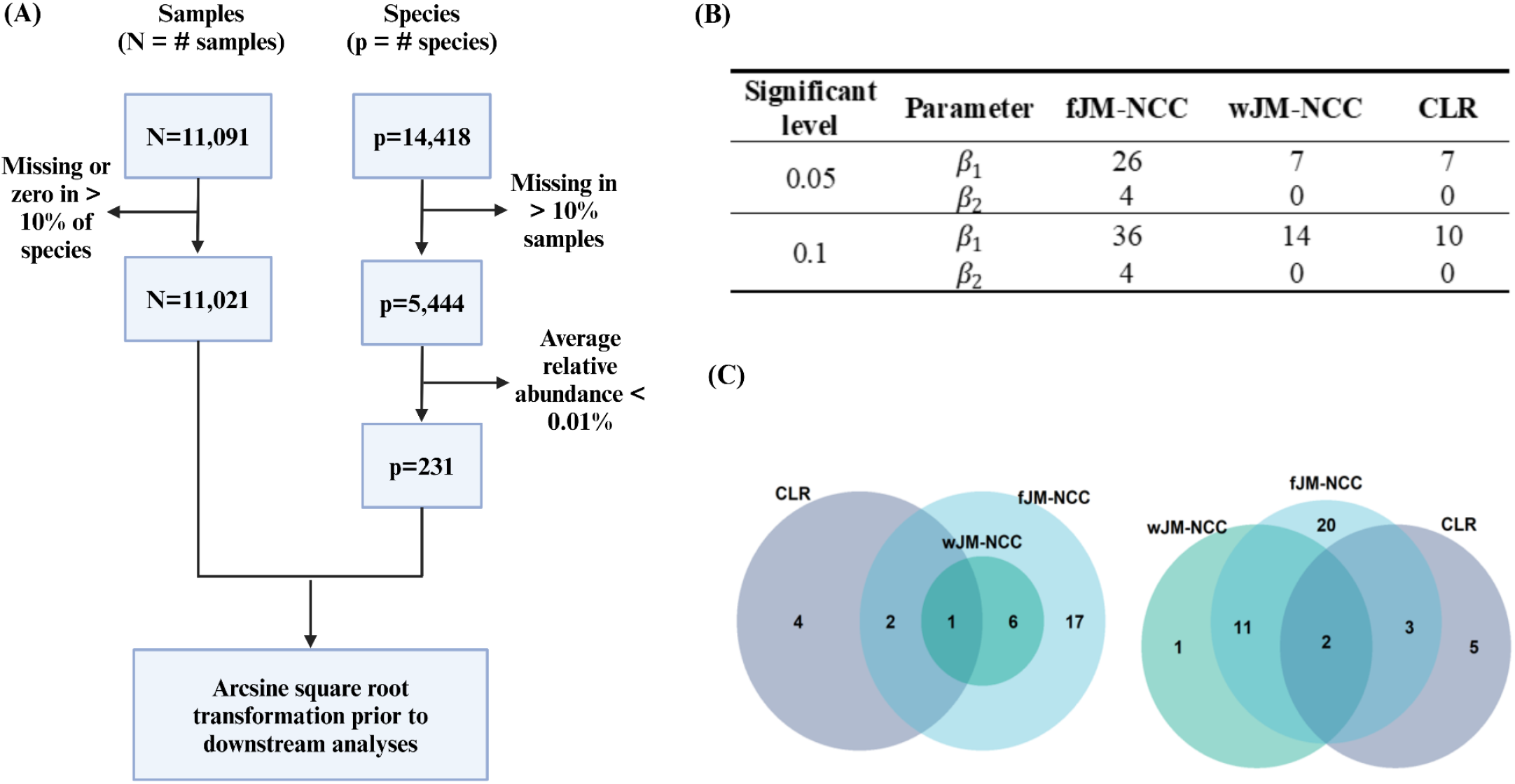
Association analysis of longitudinal microbial abundances with the competing appearance of IAA and GADA using the TEDDY Microbiome dataset. **(A):** Data filtering steps for shotgun metagenomic sequencing abundance data at the species level from the TEDDY NCC sub-cohort. **(B):** Number of significant species identified by the assessed methods (fJM-NCC, wJM-NCC, CLR, and JM) at two significant levels, 0.05 and 0.1, after Bonferroni correction. JM did not identify any significant species and is therefore excluded from the table. *β*_1_ and *β*_2_ represent the association coefficient between longitudinal species abundances and IAA-first and GADA-first respectively. **(C):** Venn diagram illustrating the overlap of significant species associated with IAA-first (*β*_1_) as identified by fJM-NCC, wJM-NCC, and CLR at nominal levels of 0.05 (left panel) and 0.1 (right panel), respectively.

**Table 5** summarizes the demographic characteristics of TEDDY cohort, stratified by their case-control status and the first appearing autoantibody. The mean event time for IAA-first was earlier than that for GADA-first, consistent with previous literature (Krischer 2015, Krischer 2017, Rewers 2018). We examined associations between each microbial biomarker (either community-level measurements or species-level relative abundances) and the competing risks of IAA-first and GADA-first by fitting association models for each biomarker and applying multiple comparison adjustments afterward. In the joint modeling analysis using fJM-NCC, wJM-NCC, and JM, birth mode and sampling time were included as fixed effects in the longitudinal sub-model, with a random intercept assumed. The longitudinal microbial trajectory, sex, birth weight, and FDR were included as covariates in the survival sub-model. Since the CLR method cannot handle competing events or time-varying covariates (microbial biomarker herein), mean community-level measures (or the mean relative abundance overtime), birth mode, sex, birth weight, and FDR of IAA-first cases and their controls, as well as those from GADA-first cases and their controls, were included in two separate models to examine microbial association with the appearance of autoantibodies.

**Table 5:** Descriptive characteristics of the TEDDY cohort. Frequency and proportions, and mean and standard deviation are summarized for categorical and continuous variables, respectively.

|  | **IAA-first** | **IAA-Control<sup>3</sup>** | **GADA-first** | **GADA-Control<sup>4</sup>** | **Controls<sup>5</sup>** | **Overall** |
| --- | --- | --- | --- | --- | --- | --- |
|  | N=244 | N=225 | N=113 | N=102 | N=7923 | N=8607 |
| Sex (Female) | 104 (42.6%) | 96 (42.7%) | 58 (51.3%) | 51 (50.0%) | 3947 (49.8%) | 4256 (49.4%) |
| FDR1 (Yes) | 54 (22.1%) | 43 (19.1%) | 19 (16.8%) | 18 (17.6%) | 801 (10.1%) | 935 (10.9%) |
| Birth Mode |  |  |  |  |  |  |
| caesarian | 51 (20.9%) | 65 (28.9%) | 28 (24.8%) | 24 (23.5%) | 2134 (26.9%) | 2302 (26.7%) |
| vaginal | 193 (79.1%) | 160 (71.1%) | 85 (75.2%) | 78 (76.5%) | 5789 (73.1%) | 6305 (73.3%) |
| Birth Weight (kg) | 3.58 (0.513) | 3.53 (0.593) | 3.62 (0.462) | 3.52 (0.541) | 3.39 (0.578) | 3.41 (0.577) |
| Event Time2 (Year) | 1.65 (1.04) | 3.99 (0.195) | 2.26 (1.32) | 4.00 (0.10) | 4.00 (0.00) | 3.91 (0.501) |

1. FDR: indicates whether any first-degree relatives in the family have T1D 2. Event Time: the time to IA onset (including IAA-first, GADA-first) or censoring time 3. IAA-Control: controls matched to IAA-first 4. GADA-Control: controls matched to GADA-first 5. Controls: controls unselected into NCC sub-cohort

**Table 6** presents the estimated parameters (*β*_1_, *β*_2_) and corresponding hypothesis testing p-values from all methods for six community-level microbiome measurements in association with the first appearance of IAA ( *β*_1_) and GADA ( *β*_2_). Results from fJM-NCC and wJM-NCC are highly consistent, as evidenced by the same direction of parameter estimates and similar levels of statistical significance. The directions of (*β*_1_, *β*_2_) estimated by fJM-NCC and wJM-NCC align well with the interpretation of competing risks. For example, both fJM-NCC and wJM-NCC indicate that lower richness and Shannon diversity (from both 16S rRNA and shotgun metagenomic sequencing data), younger microbiota age, and higher MAZ score are significantly associated with an increased risk for IAA-first. These findings are consistent with the literature suggesting that reduced microbial diversity and delayed microbial maturation are linked to poorer health outcomes and the competing nature of IAA-first and GADA-first (Krischer 2017, Stewart 2018).

**Table 6:** Estimation results and hypothesis testing p-values for the associations between six community-level microbiome measurements and appearance of IAA-first (*β*_1_) and GADA-first (*β*_2_).

| Measurements | fJM-NCC | wJM-NCC | JM | CLR |
| --- | --- | --- | --- | --- |
| *Shotgun metagenomic data:* |  |  |  |  |
| Richness (# species) | (-0.273, 0.172)<sup>1</sup><br>(0.001, 0.133)<sup>2</sup> | (-0.169, 0.229)<br>(0.026, 0.043) | (0.196, 0.006)<br>(0.171, 0.978) | (0.528, 0.435)<br>(0.004, 0.139) |
| Shannon diversity | (-0.413, 0.331)<br>(0.002, 0.075) | (-0.315, 0.370)<br>(0.010, 0.051) | (0.349, 0.399)<br>(0.182, 0.300) | (0.471, 0.463)<br>(0.098, 0.267) |
| *16S rRNA sequencing data:* |  |  |  |  |
| Richness (# OTUs) | (-0.265, 0.253)<br>($<0.001$, 0.010) | (-0.245, 0.243)<br>($<0.001$, 0.021) | (0.033, 0.390)<br>(0.850, 0.107) | (0.806, 0.638)<br>($<0.001$, 0.068) |
| Shannon diversity | (-0.415, 0.282)<br>($<0.001$, 0.071) | (-0.325, 0.322)<br>(0.002, 0.054) | (0.184, -0.074)<br>(0.450, 0.834) | (1.096, 0.931)<br>(0.001, 0.044) |
| MAZ score | (0.107, -0.087)<br>($<0.001$, 0.024) | (0.099, -0.100)<br>($<0.001$, 0.043) | (0.001, -0.001)<br>(0.984, 0.992) | (-0.093, -0.043)<br>(0.160, 0.657) |
| Microbiota Age | (-0.035, 0.028)<br>($<0.001$, 0.015) | (-0.031, 0.029)<br>($<0.001$, 0.019) | (0.030, 0.085)<br>(0.100, $<0.001$) | (0.157, 0.124)<br>($<0.001$, 0.029) |

<sup>1</sup> The point estimates and <sup>2</sup> the hypothesis testing p-values of ( $\beta_1, \beta_2$ ) for the null hypothesis $H_0: \beta_1 = 0$ and $H_0: \beta_2 = 0$ respectively.

In contrast, JM provides either identical directional estimates for *β*_1_and *β*_2_ or results entirely different from those of fJM-NCC and wJM-NCC, suggesting its limitations in NCC designed studies. CLR’s estimates (*β*_1_, *β*_2_) are in the same direction due to the model’s inability to account for the competing risks between IAA-first and GADA-first. The results of CLR indicate that higher microbial diversity, older microbiota age, and lower MAZ score are associated with higher risk of the appearance of either autoantibody, which contradicts existing findings.

The results of the species-level analysis of longitudinal microbial abundances and their association with the competing appearance of IAA-first and GADA-first are summarized in **Figure 3(B-C)** and **Table S8. Figure 3(B)** shows the number of species associated with IAA-first and GADA-first identified by each method at two nominal significant levels (0.05 and 0.1) after Bonferroni correction, while **Figure 3(C)** illustrates the overlap of species identified different methods. Table S8 provides a full list of the identified species. For IAA-first, fJM-NCC detected the highest number of significant species, identifying 26 species at the 0.05 level and 36 at the 0.1 level. For GADA-first, it identified four species at both levels. By comparison, wJM-NCC and CLR identified fewer species, with each detecting seven species for IAA-first at the 0.05 level. At the 0.1 level, wJM-NCC identified 14 species, while CLR identified 10. JM failed to identify any significant species. At the 0.05 significance level, all species identified by wJM-NCC were also detected by fJM-NCC, while CLR identified only three overlapping species with fJM-NCC. At the 0.1 level, wJM-NCC identified one species not detected by fJM-NCC, and half of the species identified by CLR overlapped with those detected by fJM-NCC. These findings demonstrate the consistency between methods, and the statistical efficiency of fJM-NCC and wJM-NCC.

We further examined two example species (**Table S8**) detected by fJM-NCC and wJM-NCC as significantly associated with autoantibody appearance through cross-referencing findings from existing biomedical literature. *Bacillus cereus* has been reported to enhance the expression of signature genes in Th17 cells, potentially delaying T1D onset (Elhag 2020). Consistent with this, the fJM-NCC estimates for *B. cereus* were (*β*_1_, *β*_2_) =(−101, −86.2) with adjusted p-values (0.019, 1), and wJM-NCC estimates were (−100, −86.1) with adjusted p-values (0.048, 1). In contrast, JM estimates were (−2.5, −0.174) with adjusted p-values (1, 1), and CLR estimates were (−1.4, 136.4) with adjusted p-values (1, 1), showing significant divergence from fJM-NCC and wJM-NCC. *Bifidobacterium breve*, a dominant gut microbiota species in infancy, has been associated with IA onset in prior studies (Li 2022). For this species, fJM-NCC estimates were (1.35, 0.454) with adjusted p-values (0.075, 0.392), while wJM-NCC estimates were (0.968, 0.131) with adjusted p-values (1, 1). However, JM estimates were (−0.092, 0.821) with adjusted p-values (1, 1), and CLR estimates were (−0.84, 0.552) with adjusted p-values (1, 1). These examples highlight the strong alignment of fJM-NCC and wJM-NCC with existing biological evidence, while JM and CLR show notable inconsistencies in their inference.

## 5 Discussion

In this paper, we propose a novel joint modeling framework specifically designed for nested case-control (NCC) studies, aimed at exploring the association between longitudinal biomarker trajectories and competing events. This framework integrates a generalized linear mixed-effects model to capture biomarker dynamics over time and a cause-specific hazard model to link these trajectories to specific competing events. We developed two maximum likelihood estimation approaches, i.e., fJM-NCC and wJM-NCC, to address the unique sampling structure and data characteristics inherent to NCC designs. fJM-NCC leverages data from both the NCC sub-cohort and the full cohort, including survival outcomes and clinical metadata. In contrast, wJM-NCC uses only NCC sub-cohort data and constructs an inverse probability weighting likelihood function to account for the potential selection bias in NCC sampling.

Simulation studies demonstrate the robustness and efficiency of both methods, as evidenced by unbiased parameter estimation and well controlled Type-I error rates across various scenarios. The statistical power of fJM-NCC is comparable to that of the Oracle method, which assumes the availability of biomarker data for the full cohort. Although wJM-NCC exhibits slightly lower power than fJM-NCC, its efficiency improves as the number of controls per case (*m*) increased, gradually approaching Oracle’s performance. In comparison, our proposed methods outperform existing approaches JM, which is unsuitable for NCC designed studies, and conditional logistic regression (CLR), which cannot effectively handle competing events and time-varying covariates.

The application of these methods to the TEDDY dataset highlights their practical utility in identifying microbial biomarkers associated with competing events, specifically IAA-first and GADA-first. The biological interpretations of six community-level microbiome measurements demonstrate the robustness of fJM-NCC and wJM-NCC in capturing meaningful associations between microbiome dynamics and disease outcomes. In the species-level microbiome association analysis, fJM-NCC identified the greatest number of significant microbial species, followed by wJM-NCC, while CLR yielded few results and JM failed to identify any.

Although this study focused on competing events within the joint modeling framework, it is important to note that the approach can also be applied to single survival outcomes as a special case when the number of competing events (*k*) is reduced to one. Many diseases, however, progress through multiple intermediate states rather than discrete competing outcomes. For example, T1D development involves transitions from the detection of autoantibodies (any IA phenotype) to the onset of overt disease (Knip 2005). Future research could extend our framework to a multi-state setting under the NCC design, enabling the investigation of biomarkers that influence disease progression across different stages. Such an extension would provide deeper insights into the dynamic roles of microbial biomarkers at various stages of disease progression, facilitating the development of more targeted and effective interventions.

## Author Contributions

YZ performed the joint modeling derivation, simulations, real data analyses, and manuscript writing. TL contributed to model derivation and simulations. BZ and CW prepared real data. AMS contributed to biological insights and interpretation. ML and HL contributed to methodology development. JH led methodology development, simulation, real data analyses, and manuscript writing.

## Acknowledgments

Not applicable.

## Supplementary Material

The supplemental material for this article can be found online.

## Data Availability Statement

The TEDDY microbiome 16S rRNA gene sequencing data and shotgun whole-genome sequencing data are publicly available in the NCBI database of Genotypes and Phenotypes (dbGaP) with the primary accession code phs001443.v1.p1, in accordance with the dbGaP controlled-access authorization process. Clinical metadata analyzed during the current study are available in the NIDDK Central Repository at https://repository.niddk.nih.gov/home.

