## Supplemental Tables and Figures for "Joint Modeling of Longitudinal Biomarker and Survival Outcomes with the Presence of Competing Risk in Nested Case-Control Studies with Application to the TEDDY Microbiome Dataset"

**Table S1:** Performance of all methods for point and 95% confidence interval estimation of $\beta_1$ and $\beta_2$ under **Scenario 2** ($\beta_1 = 0.1$ and $\beta_2 = 0.1$).

| | | $\beta_1$ | | | | | | $\beta_2$ | | | | | |
|---|---|---|---|---|---|---|---|---|---|---|---|---|---|
| **m[1]** | **Method** | **Bias** | **SE[2]** | **ESE[3]** | **MSE[4]** | **CI-L[5]** | **ECP[6]** | **Bias** | **SE** | **ESE** | **MSE** | **CI-L** | **ECP** |
| 1 | Oracle | 0.002 | 0.072 | 0.073 | 0.005 | 0.284 | 0.943 | 0.000 | 0.044 | 0.044 | 0.002 | 0.172 | 0.953 |
| | fJM-NCC | 0.003 | 0.074 | 0.077 | 0.006 | 0.292 | 0.940 | 0.001 | 0.046 | 0.048 | 0.002 | 0.182 | 0.939 |
| | wJM-NCC | 0.004 | 0.082 | 0.082 | 0.007 | 0.320 | 0.953 | 0.002 | 0.058 | 0.058 | 0.003 | 0.227 | 0.957 |
| | wJM-NCC(Fisher) | 0.004 | 0.073 | 0.082 | 0.007 | 0.284 | 0.911 | 0.002 | 0.044 | 0.058 | 0.003 | 0.172 | 0.864 |
| | JM | -0.026 | 0.072 | 0.083 | 0.008 | 0.282 | 0.933 | -0.030 | 0.044 | 0.045 | 0.003 | 0.171 | 0.899 |
| | CLR | -0.002 | 0.098 | 0.101 | 0.010 | 0.383 | 0.951 | -0.009 | 0.058 | 0.059 | 0.004 | 0.228 | 0.947 |
| 3 | Oracle | -0.003 | 0.072 | 0.073 | 0.005 | 0.284 | 0.946 | -0.003 | 0.044 | 0.043 | 0.002 | 0.172 | 0.961 |
| | fJM-NCC | -0.002 | 0.074 | 0.074 | 0.005 | 0.288 | 0.946 | -0.002 | 0.045 | 0.044 | 0.002 | 0.178 | 0.959 |
| | wJM-NCC | -0.003 | 0.075 | 0.075 | 0.006 | 0.294 | 0.943 | -0.002 | 0.049 | 0.049 | 0.002 | 0.191 | 0.946 |
| | wJM-NCC(Fisher) | -0.003 | 0.072 | 0.075 | 0.006 | 0.284 | 0.939 | -0.002 | 0.044 | 0.049 | 0.002 | 0.172 | 0.920 |
| | JM | -0.019 | 0.072 | 0.070 | 0.005 | 0.282 | 0.952 | -0.016 | 0.044 | 0.043 | 0.002 | 0.171 | 0.937 |
| | CLR | -0.007 | 0.078 | 0.079 | 0.006 | 0.305 | 0.945 | -0.011 | 0.047 | 0.046 | 0.002 | 0.184 | 0.951 |
| 5 | Oracle | 0.003 | 0.073 | 0.073 | 0.005 | 0.284 | 0.952 | 0.001 | 0.044 | 0.044 | 0.002 | 0.172 | 0.948 |
| | fJM-NCC | 0.002 | 0.073 | 0.074 | 0.005 | 0.288 | 0.952 | 0.001 | 0.045 | 0.046 | 0.002 | 0.176 | 0.940 |
| | wJM-NCC | 0.003 | 0.074 | 0.074 | 0.006 | 0.290 | 0.948 | 0.001 | 0.047 | 0.047 | 0.002 | 0.182 | 0.952 |
| | wJM-NCC(Fisher) | 0.003 | 0.072 | 0.074 | 0.006 | 0.284 | 0.945 | 0.001 | 0.044 | 0.047 | 0.002 | 0.172 | 0.930 |
| | JM | -0.007 | 0.072 | 0.073 | 0.005 | 0.282 | 0.949 | -0.008 | 0.044 | 0.044 | 0.002 | 0.171 | 0.940 |
| | CLR | -0.004 | 0.073 | 0.075 | 0.006 | 0.288 | 0.947 | -0.008 | 0.045 | 0.045 | 0.002 | 0.174 | 0.944 |

1. Control-to-case ratio, i.e., the number of controls per case in the NCC sub-cohort
2. Estimated standard error
3. Empirical standard error
4. Mean squared error
5. Average length of the 95% confidence intervals
6. Empirical coverage probability of the 95% confidence interval

**Table S2:** Performance of all methods for point and 95% confidence interval estimation of $\beta_1$ and $\beta_2$ under **Scenario 2** ($\beta_1 = 0.2$ and $\beta_2 = 0.1$).

| | | $\beta_1$ | | | | | | $\beta_2$ | | | | | |
|---|---|---|---|---|---|---|---|---|---|---|---|---|---|
| **m[1]** | **Method** | **Bias** | **SE[2]** | **ESE[3]** | **MSE[4]** | **CI-L[5]** | **ECP[6]** | **Bias** | **SE** | **ESE** | **MSE** | **CI-L** | **ECP** |
| 1 | Oracle | -0.003 | 0.071 | 0.069 | 0.005 | 0.279 | 0.963 | 0.001 | 0.044 | 0.044 | 0.002 | 0.173 | 0.941 |
| | fJM-NCC | 0.000 | 0.074 | 0.072 | 0.005 | 0.289 | 0.967 | 0.002 | 0.047 | 0.048 | 0.002 | 0.183 | 0.944 |
| | wJM-NCC | -0.002 | 0.082 | 0.080 | 0.006 | 0.320 | 0.957 | 0.001 | 0.058 | 0.059 | 0.003 | 0.227 | 0.947 |
| | wJM-NCC(Fisher) | -0.002 | 0.071 | 0.080 | 0.006 | 0.280 | 0.927 | 0.001 | 0.044 | 0.059 | 0.003 | 0.173 | 0.866 |
| | JM | -0.044 | 0.071 | 0.070 | 0.007 | 0.277 | 0.903 | -0.038 | 0.044 | 0.044 | 0.003 | 0.172 | 0.866 |
| | CLR | -0.019 | 0.098 | 0.098 | 0.010 | 0.385 | 0.953 | -0.006 | 0.059 | 0.059 | 0.004 | 0.230 | 0.945 |
| 3 | Oracle | 0.001 | 0.072 | 0.071 | 0.005 | 0.281 | 0.952 | 0.001 | 0.044 | 0.044 | 0.002 | 0.173 | 0.947 |
| | fJM-NCC | 0.003 | 0.073 | 0.073 | 0.005 | 0.286 | 0.946 | 0.002 | 0.046 | 0.045 | 0.002 | 0.179 | 0.956 |
| | wJM-NCC | 0.001 | 0.075 | 0.075 | 0.006 | 0.294 | 0.956 | 0.001 | 0.049 | 0.049 | 0.002 | 0.192 | 0.947 |
| | wJM-NCC(Fisher) | 0.001 | 0.072 | 0.075 | 0.006 | 0.281 | 0.944 | 0.001 | 0.044 | 0.049 | 0.002 | 0.173 | 0.915 |
| | JM | -0.019 | 0.071 | 0.071 | 0.005 | 0.278 | 0.952 | -0.018 | 0.044 | 0.044 | 0.002 | 0.172 | 0.928 |
| | CLR | -0.016 | 0.078 | 0.077 | 0.006 | 0.305 | 0.952 | -0.007 | 0.047 | 0.048 | 0.002 | 0.185 | 0.946 |
| 5 | Oracle | 0.001 | 0.072 | 0.077 | 0.006 | 0.281 | 0.934 | 0.003 | 0.044 | 0.046 | 0.002 | 0.173 | 0.941 |
| | fJM-NCC | 0.002 | 0.073 | 0.077 | 0.006 | 0.285 | 0.938 | 0.003 | 0.045 | 0.047 | 0.002 | 0.177 | 0.939 |
| | wJM-NCC | 0.001 | 0.073 | 0.078 | 0.006 | 0.287 | 0.945 | 0.003 | 0.047 | 0.049 | 0.002 | 0.183 | 0.942 |
| | wJM-NCC(Fisher) | 0.001 | 0.072 | 0.078 | 0.006 | 0.280 | 0.932 | 0.003 | 0.044 | 0.049 | 0.002 | 0.173 | 0.928 |
| | JM | -0.013 | 0.071 | 0.075 | 0.006 | 0.278 | 0.930 | -0.009 | 0.044 | 0.045 | 0.002 | 0.171 | 0.934 |
| | CLR | -0.017 | 0.073 | 0.079 | 0.006 | 0.287 | 0.923 | -0.005 | 0.045 | 0.046 | 0.002 | 0.176 | 0.942 |

1. Control-to-case ratio, i.e., the number of controls per case in the NCC sub-cohort
2. Estimated standard error
3. Empirical standard error
4. Mean squared error
5. Average length of the 95% confidence intervals
6. Empirical coverage probability of the 95% confidence interval

**Table S3**: Performance of all methods for point estimation of additional parameters under **Scenario 1**($\beta_1 = \beta_2 = 0.$). Parameters include the fixed slope $\gamma$, standard deviation (log) of the random intercept $\theta$, standard deviation (log) of random error $\sigma$, and the fixed effect $\alpha$.

| | | $\gamma = 0.1$ | | | $\log(\theta) = \log\sqrt{2}$ | | | $\log(\sigma) = 0$ | | | $\alpha = -0.2$ | | |
|---|---|---|---|---|---|---|---|---|---|---|---|---|---|
| **m[1]** | **Method** | **Bias** | **SE[2]** | **ESE[3]** | **Bias** | **SE** | **ESE** | **Bias** | **SE** | **ESE** | **Bias** | **SE** | **ESE** |
| 1 | Oracle | 0.000 | 0.033 | 0.033 | -0.005 | 0.008 | 0.009 | 0.001 | 0.004 | 0.004 | -0.004 | 0.101 | 0.100 |
| | fJM-NCC | 0.003 | 0.111 | 0.109 | -0.008 | 0.027 | 0.029 | 0.001 | 0.013 | 0.013 | -0.004 | 0.101 | 0.100 |
| | wJM-NCC | 0.005 | 0.144 | 0.138 | -0.007 | 0.038 | 0.039 | 0.000 | 0.017 | 0.017 | -0.004 | 0.141 | 0.103 |
| | wJM-NCC(Fisher) | 0.005 | 0.033 | 0.138 | -0.007 | 0.008 | 0.039 | 0.000 | 0.004 | 0.017 | -0.004 | 0.101 | 0.103 |
| | JM | 0.004 | 0.120 | 0.115 | 0.007 | / | 0.031 | 0.024 | / | 0.012 | 0.194 | 0.101 | 0.026 |
| 3 | Oracle | 0.000 | 0.033 | 0.032 | -0.005 | 0.008 | 0.009 | 0.001 | 0.004 | 0.004 | -0.001 | 0.101 | 0.103 |
| | fJM-NCC | -0.002 | 0.078 | 0.082 | -0.005 | 0.020 | 0.020 | 0.001 | 0.009 | 0.009 | -0.001 | 0.101 | 0.103 |
| | wJM-NCC | -0.004 | 0.086 | 0.086 | -0.005 | 0.022 | 0.022 | 0.001 | 0.010 | 0.010 | 0.000 | 0.115 | 0.105 |
| | wJM-NCC(Fisher) | -0.004 | 0.033 | 0.086 | -0.005 | 0.008 | 0.022 | 0.001 | 0.004 | 0.010 | 0.000 | 0.101 | 0.105 |
| | JM | -0.002 | 0.086 | 0.088 | 0.009 | / | 0.021 | 0.025 | / | 0.009 | 0.184 | 0.101 | 0.019 |
| 5 | Oracle | 0.000 | 0.033 | 0.033 | -0.005 | 0.008 | 0.009 | 0.001 | 0.004 | 0.004 | 0.000 | 0.101 | 0.100 |
| | fJM-NCC | -0.001 | 0.064 | 0.067 | -0.006 | 0.016 | 0.017 | 0.001 | 0.008 | 0.008 | 0.000 | 0.101 | 0.100 |
| | wJM-NCC | 0.001 | 0.068 | 0.070 | -0.006 | 0.018 | 0.018 | 0.001 | 0.008 | 0.008 | 0.000 | 0.109 | 0.101 |
| | wJM-NCC(Fisher) | 0.001 | 0.033 | 0.070 | -0.006 | 0.008 | 0.018 | 0.001 | 0.004 | 0.008 | 0.000 | 0.101 | 0.101 |
| | JM | -0.001 | 0.071 | 0.069 | 0.009 | / | 0.018 | 0.025 | / | 0.007 | 0.173 | 0.101 | 0.020 |

1. Control-to-case ratio, i.e., the number of controls per case in the NCC sub-cohort
2. Estimated standard error
3. Empirical standard error

**Table S4**: Performance of all methods for point estimation of additional parameters under **Scenario 2**($\beta_1 = 0$ and $\beta_2 = 0.1$). Parameters include the fixed slope γ, standard deviation (log) of the random intercept $\theta$, standard deviation (log) of random error $\sigma$, and the fixed effect α.

| | | $\gamma = 0.1$ | | | $\log(\theta) = \log\sqrt{2}$ | | | $\log(\sigma) = 0$ | | | $\alpha = -0.2$ | | |
|---|---|---|---|---|---|---|---|---|---|---|---|---|---|
| **m**[1] | **Method** | **Bias** | **SE**[2] | **ESE**[3] | **Bias** | **SE** | **ESE** | **Bias** | **SE** | **ESE** | **Bias** | **SE** | **ESE** |
| 1 | Oracle | -0.001 | 0.033 | 0.034 | -0.005 | 0.008 | 0.009 | 0.001 | 0.004 | 0.004 | -0.002 | 0.101 | 0.098 |
| | fJM-NCC | -0.002 | 0.111 | 0.113 | -0.007 | 0.027 | 0.028 | 0.000 | 0.013 | 0.013 | -0.002 | 0.101 | 0.098 |
| | wJM-NCC | -0.002 | 0.144 | 0.136 | -0.008 | 0.038 | 0.039 | 0.001 | 0.017 | 0.018 | -0.002 | 0.141 | 0.100 |
| | wJM-NCC(Fisher) | -0.002 | 0.033 | 0.136 | -0.008 | 0.008 | 0.039 | 0.001 | 0.004 | 0.018 | -0.002 | 0.101 | 0.100 |
| | JM | 0.001 | 0.120 | 0.122 | 0.010 | / | 0.029 | 0.024 | / | 0.013 | 0.194 | 0.101 | 0.027 |
| 3 | Oracle | 0.001 | 0.033 | 0.032 | -0.005 | 0.008 | 0.009 | 0.001 | 0.004 | 0.004 | -0.002 | 0.101 | 0.101 |
| | fJM-NCC | -0.002 | 0.078 | 0.077 | -0.007 | 0.020 | 0.020 | 0.001 | 0.009 | 0.009 | -0.002 | 0.101 | 0.101 |
| | wJM-NCC | -0.005 | 0.086 | 0.084 | -0.007 | 0.022 | 0.022 | 0.001 | 0.010 | 0.010 | -0.002 | 0.115 | 0.102 |
| | wJM-NCC(Fisher) | -0.005 | 0.033 | 0.084 | -0.007 | 0.008 | 0.022 | 0.001 | 0.004 | 0.010 | -0.002 | 0.101 | 0.102 |
| | JM | -0.005 | 0.086 | 0.082 | 0.009 | / | 0.022 | 0.025 | / | 0.009 | 0.182 | 0.101 | 0.019 |
| 5 | Oracle | 0.001 | 0.033 | 0.033 | -0.005 | 0.008 | 0.009 | 0.001 | 0.004 | 0.004 | 0.002 | 0.101 | 0.097 |
| | fJM-NCC | 0.000 | 0.064 | 0.063 | -0.006 | 0.016 | 0.017 | 0.001 | 0.008 | 0.008 | 0.002 | 0.101 | 0.097 |
| | wJM-NCC | 0.000 | 0.068 | 0.067 | -0.005 | 0.018 | 0.017 | 0.001 | 0.008 | 0.008 | 0.002 | 0.109 | 0.098 |
| | wJM-NCC(Fisher) | 0.000 | 0.033 | 0.067 | -0.005 | 0.008 | 0.017 | 0.001 | 0.004 | 0.008 | 0.002 | 0.101 | 0.098 |
| | JM | 0.000 | 0.071 | 0.068 | 0.010 | / | 0.018 | 0.025 | / | 0.007 | 0.173 | 0.101 | 0.021 |

1. Control-to-case ratio, i.e., the number of controls per case in the NCC sub-cohort
2. Estimated standard error
3. Empirical standard error

**Table S5**: Performance of all methods for point estimation of additional parameters under **Scenario 2**($\beta_1 = 0.1$ and $\beta_2 = 0.1$). Parameters include the fixed slope $\gamma$, standard deviation (log) of the random intercept $\theta$, standard deviation (log) of random error $\sigma$, and the fixed effect $\alpha$.

| | | $\gamma = 0.1$ | | | $\log(\theta) = \log\sqrt{2}$ | | | $\log(\sigma) = 0$ | | | $\alpha = -0.2$ | | |
|---|---|---|---|---|---|---|---|---|---|---|---|---|---|
| **m[1]** | **Method** | **Bias** | **SE[2]** | **ESE[3]** | **Bias** | **SE** | **ESE** | **Bias** | **SE** | **ESE** | **Bias** | **SE** | **ESE** |
| 1 | Oracle | 0.001 | 0.033 | 0.034 | -0.004 | 0.008 | 0.009 | 0.001 | 0.004 | 0.004 | 0.004 | 0.101 | 0.102 |
| | fJM-NCC | 0.001 | 0.111 | 0.112 | -0.006 | 0.027 | 0.029 | 0.000 | 0.013 | 0.013 | 0.004 | 0.101 | 0.102 |
| | wJM-NCC | 0.001 | 0.144 | 0.141 | -0.005 | 0.038 | 0.038 | 0.001 | 0.017 | 0.018 | 0.005 | 0.142 | 0.105 |
| | wJM-NCC(Fisher) | 0.001 | 0.033 | 0.141 | -0.005 | 0.008 | 0.038 | 0.001 | 0.004 | 0.018 | 0.005 | 0.101 | 0.105 |
| | JM | 0.004 | 0.120 | 0.120 | 0.011 | / | 0.033 | 0.024 | / | 0.013 | 0.193 | 0.101 | 0.042 |
| 3 | Oracle | -0.001 | 0.033 | 0.032 | -0.005 | 0.008 | 0.009 | 0.001 | 0.004 | 0.004 | 0.001 | 0.101 | 0.101 |
| | fJM-NCC | -0.001 | 0.078 | 0.080 | -0.005 | 0.020 | 0.020 | 0.002 | 0.009 | 0.010 | 0.001 | 0.101 | 0.101 |
| | wJM-NCC | -0.004 | 0.086 | 0.088 | -0.005 | 0.022 | 0.022 | 0.002 | 0.010 | 0.011 | 0.001 | 0.115 | 0.102 |
| | wJM-NCC(Fisher) | -0.004 | 0.033 | 0.088 | -0.005 | 0.008 | 0.022 | 0.002 | 0.004 | 0.011 | 0.001 | 0.101 | 0.102 |
| | JM | -0.003 | 0.086 | 0.086 | 0.011 | / | 0.022 | 0.026 | / | 0.009 | 0.182 | 0.101 | 0.020 |
| 5 | Oracle | 0.000 | 0.033 | 0.032 | -0.005 | 0.008 | 0.009 | 0.001 | 0.004 | 0.004 | 0.000 | 0.101 | 0.104 |
| | fJM-NCC | 0.004 | 0.064 | 0.067 | -0.005 | 0.016 | 0.017 | 0.001 | 0.008 | 0.008 | 0.000 | 0.101 | 0.104 |
| | wJM-NCC | 0.003 | 0.068 | 0.070 | -0.004 | 0.018 | 0.018 | 0.001 | 0.008 | 0.008 | 0.000 | 0.109 | 0.106 |
| | wJM-NCC(Fisher) | 0.003 | 0.033 | 0.070 | -0.004 | 0.008 | 0.018 | 0.001 | 0.004 | 0.008 | 0.000 | 0.101 | 0.106 |
| | JM | 0.004 | 0.071 | 0.071 | 0.011 | / | 0.018 | 0.026 | / | 0.007 | 0.173 | 0.101 | 0.022 |

1. Control-to-case ratio, i.e., the number of controls per case in the NCC sub-cohort
2. Estimated standard error
3. Empirical standard error

**Table S6**: Performance of all methods for point estimation of additional parameters under **Scenario 2**($\beta_1 = 0.2$ and $\beta_2 = 0.1$). Parameters include the fixed slope $\gamma$, standard deviation (log) of the random intercept $\theta$, standard deviation (log) of random error $\sigma$, and the fixed effect α.

| | | $\gamma = 0.1$ | | | $\log(\theta) = \log\sqrt{2}$ | | | $\log(\sigma) = 0$ | | | $\alpha = -0.2$ | | |
|---|---|---|---|---|---|---|---|---|---|---|---|---|---|
| **m[1]** | **Method** | **Bias** | **SE[2]** | **ESE[3]** | **Bias** | **SE** | **ESE** | **Bias** | **SE** | **ESE** | **Bias** | **SE** | **ESE** |
| 1 | Oracle | 0.001 | 0.033 | 0.032 | -0.005 | 0.008 | 0.009 | 0.001 | 0.004 | 0.004 | -0.004 | 0.101 | 0.098 |
| | fJM-NCC | -0.001 | 0.110 | 0.112 | -0.006 | 0.027 | 0.028 | 0.001 | 0.013 | 0.013 | -0.005 | 0.101 | 0.098 |
| | wJM-NCC | -0.003 | 0.144 | 0.145 | -0.004 | 0.038 | 0.039 | 0.000 | 0.017 | 0.018 | -0.006 | 0.142 | 0.102 |
| | wJM-NCC(Fisher) | -0.003 | 0.033 | 0.145 | -0.004 | 0.008 | 0.039 | 0.000 | 0.004 | 0.018 | -0.006 | 0.101 | 0.102 |
| | JM | -0.001 | 0.121 | 0.119 | 0.014 | / | 0.031 | 0.025 | / | 0.012 | 0.190 | 0.101 | 0.028 |
| 3 | Oracle | -0.001 | 0.033 | 0.033 | -0.005 | 0.008 | 0.009 | 0.001 | 0.004 | 0.004 | 0.001 | 0.101 | 0.100 |
| | fJM-NCC | 0.003 | 0.077 | 0.078 | -0.006 | 0.020 | 0.020 | 0.001 | 0.009 | 0.009 | 0.001 | 0.101 | 0.100 |
| | wJM-NCC | 0.001 | 0.085 | 0.086 | -0.006 | 0.022 | 0.023 | 0.001 | 0.010 | 0.010 | 0.002 | 0.115 | 0.102 |
| | wJM-NCC(Fisher) | 0.001 | 0.033 | 0.086 | -0.006 | 0.008 | 0.023 | 0.001 | 0.004 | 0.010 | 0.002 | 0.101 | 0.102 |
| | JM | 0.002 | 0.086 | 0.084 | 0.012 | / | 0.021 | 0.025 | / | 0.009 | 0.183 | 0.101 | 0.021 |
| 5 | Oracle | -0.001 | 0.033 | 0.034 | -0.005 | 0.008 | 0.009 | 0.001 | 0.004 | 0.004 | 0.000 | 0.101 | 0.100 |
| | fJM-NCC | 0.000 | 0.064 | 0.065 | -0.005 | 0.016 | 0.016 | 0.001 | 0.008 | 0.007 | 0.000 | 0.101 | 0.100 |
| | wJM-NCC | 0.000 | 0.068 | 0.068 | -0.004 | 0.018 | 0.017 | 0.001 | 0.008 | 0.008 | 0.000 | 0.109 | 0.102 |
| | wJM-NCC(Fisher) | 0.000 | 0.033 | 0.068 | -0.004 | 0.008 | 0.017 | 0.001 | 0.004 | 0.008 | 0.000 | 0.101 | 0.102 |
| | JM | 0.000 | 0.071 | 0.071 | 0.012 | / | 0.018 | 0.026 | / | 0.007 | 0.173 | 0.101 | 0.022 |

1. Control-to-case ratio, i.e., the number of controls per case in the NCC sub-cohort
2. Estimated standard error
3. Empirical standard error

**Table S7**: Performance of all methods for point estimation of additional parameters under **Scenario 2**($\beta_1 = 0.3$ and $\beta_2 = 0.1$). Parameters include the fixed slope $\gamma$, standard deviation (log) of the random intercept $\theta$, standard deviation (log) of random error $\sigma$, and the fixed effect $\alpha$.

| | | $\gamma = 0.1$ | | | $\log(\theta) = \log\sqrt{2}$ | | | $\log(\sigma) = 0$ | | | $\alpha = -0.2$ | | |
|---|---|---|---|---|---|---|---|---|---|---|---|---|---|
| **m[1]** | **Method** | **Bias** | **SE[2]** | **ESE[3]** | **Bias** | **SE** | **ESE** | **Bias** | **SE** | **ESE** | **Bias** | **SE** | **ESE** |
| 1 | Oracle | 0.000 | 0.033 | 0.033 | -0.004 | 0.008 | 0.008 | 0.001 | 0.004 | 0.004 | -0.004 | 0.101 | 0.097 |
| | fJM-NCC | 0.007 | 0.108 | 0.104 | -0.006 | 0.027 | 0.027 | 0.000 | 0.013 | 0.012 | -0.004 | 0.101 | 0.097 |
| | wJM-NCC | 0.004 | 0.144 | 0.137 | -0.005 | 0.038 | 0.038 | 0.000 | 0.017 | 0.016 | -0.003 | 0.143 | 0.101 |
| | wJM-NCC(Fisher) | 0.004 | 0.033 | 0.137 | -0.005 | 0.008 | 0.038 | 0.000 | 0.004 | 0.016 | -0.003 | 0.101 | 0.101 |
| | JM | 0.008 | 0.121 | 0.113 | 0.019 | / | 0.029 | 0.025 | / | 0.012 | 0.191 | 0.101 | 0.089 |
| 3 | Oracle | 0.001 | 0.033 | 0.031 | -0.004 | 0.008 | 0.009 | 0.001 | 0.004 | 0.004 | 0.000 | 0.101 | 0.100 |
| | fJM-NCC | 0.001 | 0.077 | 0.076 | -0.005 | 0.020 | 0.020 | 0.001 | 0.009 | 0.009 | 0.000 | 0.101 | 0.100 |
| | wJM-NCC | 0.002 | 0.085 | 0.084 | -0.005 | 0.022 | 0.022 | 0.001 | 0.010 | 0.010 | 0.000 | 0.116 | 0.102 |
| | wJM-NCC(Fisher) | 0.002 | 0.033 | 0.084 | -0.005 | 0.008 | 0.022 | 0.001 | 0.004 | 0.010 | 0.000 | 0.101 | 0.102 |
| | JM | 0.000 | 0.086 | 0.083 | 0.016 | / | 0.021 | 0.026 | / | 0.009 | 0.182 | 0.101 | 0.023 |
| 5 | Oracle | 0.000 | 0.033 | 0.033 | -0.005 | 0.008 | 0.009 | 0.001 | 0.004 | 0.004 | 0.001 | 0.101 | 0.102 |
| | fJM-NCC | 0.002 | 0.064 | 0.064 | -0.005 | 0.016 | 0.016 | 0.000 | 0.008 | 0.008 | 0.001 | 0.101 | 0.102 |
| | wJM-NCC | 0.000 | 0.068 | 0.068 | -0.005 | 0.018 | 0.017 | 0.000 | 0.008 | 0.008 | 0.002 | 0.109 | 0.106 |
| | wJM-NCC(Fisher) | 0.000 | 0.033 | 0.068 | -0.005 | 0.008 | 0.017 | 0.000 | 0.004 | 0.008 | 0.002 | 0.101 | 0.106 |
| | JM | 0.002 | 0.071 | 0.070 | 0.014 | / | 0.018 | 0.025 | / | 0.007 | 0.174 | 0.101 | 0.024 |

1. Control-to-case ratio, i.e., the number of controls per case in the NCC sub-cohort
2. Estimated standard error
3. Empirical standard error

**Table S8:** Estimation results and hypothesis testing p-values for the species that associated with appearance of IAA-first ($\beta_1$) and GADA-first ($\beta_2$) identified by various methods.

| Species | fJM-NCC | wJM-NCC | CLR |
|---|---|---|---|
| *Megasphaera elsdenii* | (-40.50, -48.40)[1]<br>(< 0.001, < 0.001)[2] | (-38.90, -46.40)<br>(1, 1) | (8.97, 15.64)<br>(1, 1) |
| *Bifidobacterium dentium* | (4.75, 4.68)<br>(< 0.001, < 0.001) | (-5.98, -5.05)<br>(0.199, 1.00) | (-10.66, -2.26)<br>(0.639, 1) |
| *Lactobacillus delbrueckii* | (-80.20, 22.70)<br>(0.001, 1) | (-58.80, 42.20)<br>(0.322, 1) | (67.25, -11.26)<br>(1, 1) |
| *Bacteroides heparinolyticus* | (-35.80, 5.20)<br>(0.001, 1) | (-35.70, 5.25)<br>(0.040, 1) | (11.06, 14.79)<br>(1, 1) |
| *Ethanoligenens harbinense* | (-43.3, 26.7)<br>(0.002, 1) | (-30.9, 30.7)<br>(0.138, 1) | (105.518, 87.925)<br>(0.153, 1) |
| *Acidaminococcus intestini* | (-36.7, -16)<br>(0.002, 1) | (-36.1, -15.2)<br>(1, 1) | (22.051, -5.407)<br>(1, 1) |
| *Bacteroides sp. A1C1* | (-7, 0.412)<br>(0.003, 1) | (-5.59, 1.1)<br>(0.082, 1) | (-1.161, -3.188)<br>(1, 1) |
| *Escherichia marmotae* | (116, -52.6)<br>(0.004, 1) | (116, -52.8)<br>(0.118, 1) | (23.832, -8.44)<br>(1, 1) |
| *Oscillibacter valericigenes* | (-31.9, 24.4)<br>(0.012, 1) | (-26.4, 24.6)<br>(0.060, 1) | (78.761, 46.646)<br>(0.401, 1) |
| *Corynebacterium argentoratense* | (-50.8, -2.79)<br>(0.013, 1) | (-50.5, -2.53)<br>(0.225, 1) | (10.472, -15.795)<br>(1, 1) |
| *Gordonibacter urolithinfaciens* | (-34.8, 10.7)<br>(0.013, 1) | (-34.7, 10.6)<br>(0.050, 1) | (4.748, -4.489)<br>(1, 1) |
| *Ruminococcus albus* | (-31.6, 23.6)<br>(0.014, 1) | (-23.2, 27.2)<br>(0.543, 1) | (98.702, 69.7)<br>(0.028, 1) |
| *Ruminococcaceae bacterium CPB6* | (-34, 27.8)<br>(0.014, 1) | (-26.2, 29.6)<br>(0.242, 1) | (107.033, 79.485)<br>(0.024, 1) |
| *Caproiciproducens sp. NJN-50* | (-35, 30.4)<br>(0.014, 1) | (-28.9, 31.3)<br>(0.091, 1) | (91.71, 89.276)<br>(0.138, 1) |
| *Paraprevotella xylaniphila* | (-29.7, 17.6)<br>(0.015, 1) | (-28.7, 17.9)<br>(0.116, 1) | (14.925, 10.281)<br>(1, 1) |
| *Bacillus cereus* | (-101, -86.2)<br>(0.019, 1) | (-100, -86.1)<br>(0.048, 1) | (-1.47, 136.413)<br>(1, 1) |
| *Bacteroides intestinalis* | (-18.4, 0.944)<br>(0.024, 1) | (-10.9, 6)<br>(1, 1) | (10.539, 7.884)<br>(1, 1) |
| *Escherichia albertii* | (77.2, -41.4)<br>(0.026, 1) | (56.4, -54.7)<br>(1, 1) | (-0.466, -78.615)<br>(1, 1) |
| *Alistipes finegoldii* | (-7.84, 4.15)<br>(0.029, 1) | (-7.29, 4.35)<br>(0.061, 1) | (2.133, 0.458)<br>(1, 1) |
| *Ruminococcus bicirculans* | (-6.59, 6.16)<br>(0.033, 1) | (-5.63, 7.24)<br>(0.281, 1) | (14.039, 4.357)<br>(0.729, 1) |
| *Christensenella sp. Marseille-P3954* | (-32.3, 23.9)<br>(0.033, 1) | (-32, 23.9)<br>(0.012, 1) | (88.909, 62.8)<br>(0.168, 1) |

| | | | |
|---|---|---|---|
| *Streptococcus thermophilus* | (-16.8, 2.91)<br>(0.033, 1) | (-16.5, 2.77)<br>(0.088, 1) | (0.081, -3.191)<br>(1, 1) |
| *Corynebacterium variabile* | (-314, -362)<br>(0.033, 0.315) | (-314, -362)<br>(0.003, 0.375) | (57.346, -97.577)<br>(1, 1) |
| *Ruminococcus champanellensis* | (-23.3, 22.1)<br>(0.038, 1) | (-23.1, 22.2)<br>(0.027, 1) | (77.238, 34.679)<br>(0.040, 1) |
| *Alistipes sp. 5CBH24* | (-29, 26.6)<br>(0.043, 1) | (-29, 26.4)<br>(0.008, 1) | (19.585, 7.32)<br>(1, 1) |
| *Flintibacter sp. KGMB00164* | (-18.7, 13.8)<br>(0.045, 1) | (-15.9, 13.9)<br>(0.156, 1) | (50.723, 18.737)<br>(0.282, 1) |
| *Bifidobacterium* | (-1.3, 0.059)<br>(0.059, 1) | (0.491, 1.76)<br>(1, 1) | (2.064, -0.768)<br>(1, 1) |
| *Clostridiales bacterium CCNA10* | (-10.2, 7.09)<br>(0.067, 1) | (-8.69, 7.02)<br>(0.132, 1) | (31.273, 17.425)<br>(0.087, 1) |
| *Ruminococcus sp. JE7A12* | (-11.1, 9.24)<br>(0.073, 1) | (-8.77, 11.7)<br>(1, 1) | (25.473, 20.154)<br>(0.591, 1) |
| *Veillonella parvula* | (6.63, -5.88)<br>(0.075, 1) | (6.08, -6.16)<br>(0.354, 1) | (-3.582, -4.292)<br>(1, 1) |
| *Bifidobacterium breve* | (1.35, 0.454)<br>(0.075, 1) | (0.968, 0.131)<br>(1, 1) | (-0.84, 0.552)<br>(1, 1) |
| *Bacteroides dorei* | (-3.41, -1.62)<br>(0.075, 1) | (-2.04, -0.081)<br>(1, 1) | (-0.878, 1.489)<br>(1, 1) |
| *Intestinimonas butyriciproducens* | (-25.9, 21)<br>(0.084, 1) | (-25.6, 21.2)<br>(0.083, 1) | (93.448, 37.092)<br>(0.039, 1) |
| *Adlercreutzia equolifaciens* | (-18.5, 11.8)<br>(0.090, 1) | (-18.4, 11.8)<br>(0.208, 1) | (31.881, 10.543)<br>(1, 1) |
| *Paeniclostridium sordellii* | (-43.9, 19.9)<br>(0.093, 1) | (-21.9, 28.6)<br>(1, 1) | (40.45, 70.657)<br>(1, 1) |
| *Roseburia hominis* | (-7.71, 6)<br>(0.095, 1) | (-5.89, 6.82)<br>(1, 1) | (16.917, 18.309)<br>(0.596, 1) |
| *Collinsella aerofaciens* | (2.85, 7.19)<br>(0.276, <0.001) | (-0.603, 3.86)<br>(1, 0.789) | (0.395, 5.9)<br>(1, 1) |
| *Lactobacillus paracasei* | (-11.4, -22.3)<br>(0.242, 0.002) | (3.26, -24.7)<br>(1, 1) | (-7.275, -19.401)<br>(1, 1) |
| *Alistipes sp. 5CPEGH6* | (-23.8, 20)<br>(0.170, 1) | (-23.6, 19.9)<br>(0.098, 1) | (50.942, 3.761)<br>(1, 1) |
| *[Eubacterium] sulci* | (-29.8, 12.2)<br>(1, 1) | (-39.7, 61.1)<br>(1, 1) | (167.44, 122.065)<br>(0.009, 1) |
| *Lachnospiraceae bacterium GAM79* | (-6.4, 5.56)<br>(0.134, 1) | (-4.76, 6.91)<br>(1, 1) | (19.413, 10.612)<br>(0.020, 1) |
| *Turicibacter sp. H121* | (-15.9, 14.2)<br>(1, 1) | (-10.4, 17.2)<br>(1, 1) | (48.578, 32.014)<br>(0.048, 1) |
| *Anaerobutyricum hallii* | (-5.14, 4.31)<br>(0.566, 1) | (-3.79, 5.02)<br>(1, 1) | (17.125, 13.296)<br>(0.062, 1) |
| *Campylobacter jejuni* | (-13.4, 0.894)<br>(1, 1) | (-13.4, 1.1)<br>(1, 1) | (90.038, -28.752)<br>(0.062, 1) |

[1] The point estimates and [2] the hypothesis testing p-values of $(\beta_1, \beta_2)$ for the null hypothesis $H_0: \beta_1 = 0$ and $H_0: \beta_2 = 0$ respectively.
